# Optimal Closed-loop Deep Brain Stimulation with Multi-Contact Electrodes

**DOI:** 10.1101/2020.08.10.242743

**Authors:** Gihan Weerasinghe, Benoit Duchet, Christian Bick, Rafal Bogacz

**Affiliations:** MRC Brain Network Dynamics Unit, Nuffield Department of Clinical Neurosciences, University of Oxford, Oxford, UK; Oxford Centre for Industrial and Applied Mathematics, Mathematical Institute, University of Oxford, Oxford, UK; Centre for Systems, Dynamics and Control and Department of Mathematics, University of Exeter, Exeter, UK; EPSRC Centre for Predictive Modelling in Healthcare, University of Exeter, Exeter, UK

## Abstract

Deep brain stimulation (DBS) is a well-established treatment option for a variety of neurological disorders, including Parkinson’s disease (PD) and essential tremor (ET). It is widely believed that the efficacy, efficiency and side-effects of the treatment can be improved by stimulating ‘closed-loop’, according to the symptoms of a patient. Multi-contact electrodes powered by independent current sources are a recent development in DBS technology which allow for greater precision when targeting one or more pathological regions but, in order to realise the potential of such systems, algorithms must be developed to deal with their increased complexity. This motivates the need to understand how applying DBS to multiple regions (or neural populations) can affect the efficacy and efficiency of the treatment. On the basis of a theoretical model, our paper aims to address the question of how to best apply DBS to multiple neural populations to maximally desynchronise brain activity. Using a coupled oscillator model, we derive analytical expressions which predict how the symptom severity should change as a result of applying stimulation. On the basis of these expressions we derive an algorithm describing when the stimulation should be delivered to individual contacts. Remarkably, these expressions also allow us to determine the conditions for when stimulation using information from individual contacts is likely to be advantageous. Using numerical simulation, we demonstrate that our methods have the potential to be both more effective and efficient than existing methods found in the literature.

## 1. Introduction

Deep brain stimulation (DBS) is an effective treatment for advanced Parkinson’s disease (PD) and essential tremor (ET) which involves delivering stimulation through electrodes implanted deep into the brain and targeting regions thought to be implicated in the disease, which in the case of PD is typically the subthalamic nucleus (STN) and for ET the ventral intermediate nucleus (VIM). PD is a common movement disorder caused by the death of dopaminergic neurons in the substantia nigra. Primarily, symptoms manifest as slowness of movement (bradykinesia), muscle stiffness (rigidity) and tremor. ET is purportedly the most common movement disorder, affecting just under 1% of the world population [1, 2] with the main symptom being involuntary shaking most commonly in the upper limbs [3]. Despite its prevalence, the pathophysiology of ET remains elusive, although the cortex, thalamus and cerebellum are all thought to be involved in the disease [2]. Symptoms of these disorders are thought to be due to overly synchronous activity within neural populations. For PD patients, higher power in the beta frequency range (13-30Hz) of the local field potential (LFP) measured in the STN has been shown to correlate with motor impairment [4] while thalamic activity in ET patients is strongly correlated with tremor measured using the wrist flexor EMG [5]. It is thought that DBS acts to desynchronise this pathological activity leading to a reduction in the symptom severity.

A typical DBS system consists of a lead, an implantable pulse generator (IPG) and a unit to be operated by the patient. The DBS lead terminates with an electrode, which is typically divided into multiple contacts. Post surgery, clinicians manually tune the various parameters of stimulation, such as the frequency, amplitude and pulse width, in an attempt to achieve optimal therapeutic benefit. Stimulation is then provided constantly, or ‘open-loop’, according to these parameters. The choice of stimulation frequency in particular is known to be crucial for efficacy with high frequency (HF) DBS (120-180 Hz) being found to be effective for both PD and ET patients [6]. Despite the effectiveness of conventional HF DBS in treating PD and ET, it is believed that improvements to the efficiency and efficacy can be achieved by using more elaborate stimulation patterns informed by mathematical models. Coordinated reset (CR) neuromodulation is an open-loop DBS strategy where brief HF pulse trains are applied through different contacts of a stimulation electrode [7, 8, 9, 10]. The efficacy of CR was first demonstrated theoretically, where precisely-timed delivery of HF pulses can be shown to desynchronise a system of coupled oscillators [7]. In practice, CR has been shown to yield both acute and long-lasting benefits in nonhuman primates [8].

Closed-loop stimulation and IPGs with multiple independent current sources are promising new advances in DBS technology. Closed-loop stimulation is a new development in DBS methods which aims to deliver stimulation on the basis of feedback from a patient. There is a growing body of evidence [11, 12, 4, 13] suggesting that closed-loop stimulation has the potential to offer improvements in terms of efficacy, efficiency and reduction in side effects. IPGs with multiple independent current sources are the ‘cutting-edge’ of DBS technology which, unlike their single current source counterparts, allow for current to be delivered independently to each contact. This gives increased control and flexibility over the shape of the electric fields delivered through the electrodes, allowing for more precise targeting of pathological regions and the possibility of delivering more complex potential fields over space, in addition to allowing for the possibility of recording activity from different regions. The use of multiple contacts for DBS, however, naturally leads to increased complexity, as many more stimulation strategies are now possible. This has created the need to better understand how applying DBS through multiple contacts can affect the treatment.

Closed-loop DBS strategies are characterised by their use of a feedback signal to determine when stimulation should be applied. The choice, use and accuracy of this feedback signal therefore plays a crucial role in determining the efficacy of a particular strategy. In the literature, both the LFP and tremor have been used as feedback signals with studies showing that the effects of DBS to be dependent on both the phase and amplitude of the oscillations at the time of stimulation [12, 4]. In adaptive DBS, high frequency stimulation is applied only when the amplitude of oscillations exceeds a certain threshold [4] and in phase-locked DBS stimulation is applied according to the instantaneous phase of the oscillations, which for ET patients corresponds to stimulation at roughly the tremor frequency (typically *∼* 5 Hz) [12]. The combined approach of adaptive and phase-locked stimulation has also been investigated in simulation [13].

In our previous work [14], we provided a mathematical basis for the phase and amplitude dependence of DBS. Here, we extend these ideas and introduce adaptive coordinated reset (ACR), which proposes a closed-loop strategy especially suited to multi-contact systems. Our goal is to understand how the effects of multi-contact DBS should depend on the ongoing neural activity measured through each channel. As part of this work, we demonstrate using numerical simulations that a coupled oscillator model is a plausible neural mechanism for generating tremor found in ET patients. Then, on the basis of this, we model the activity of multiple neural populations using a set of oscillators and relate this activity to the pathological oscillations associated with symptom severity in ET and PD. Using a coupled oscillator model we then describe how this activity (and hence the symptom severity) is likely to change when DBS is applied through multiple contacts. The results we present suggest how DBS should be provided through multiple contacts in order to maximally desynchronise neural activity. Using numerical simulation and parameters fitted to ET patients, we then compare our methods to others found in the literature, namely phase-locked stimulation and coordinated reset. The methods we present can be applied in different ways, either using multiple electrodes or single electrodes with multiple contacts. We therefore use the terms ‘electrode’ and ‘contact’ synonymously throughout.

## 2. A Model for Single Contact DBS

### 2.1. Phase Synchrony and Oscillations

In this section, we consider how stimulation with a single electrode acts on a population of oscillators. Here we follow our previous paper [14], which the interested reader may refer to for a more detailed derivation of the results presented in this section. A list of frequently used notation is provided in Table 1.

**Table 1.** List of frequently used symbols together with their description.

| Parameter | Description |
| --- | --- |
| $k$ | Coupling constant |
| $\tilde{\sigma}$ | Noise amplitude |
| $\bar{\omega}$ | Mean of natural frequencies |
| $s_{\omega}$ | Standard deviation of natural frequencies |
| $Z$ | Neuronal phase response curve (nPRC) |
| $a$ | Cosine Fourier coefficient of nPRC |
| $b$ | Sine Fourier coefficient of nPRC |
| $\theta$ | Phase of oscillator |
| $\psi$ | Phase of population |
| $\rho$ | Synchrony of population |
| $r$ | Complex order parameter |
| $c$ | Scaling constant for experimental data |
| $S$ | Number of populations |
| $L$ | Number of electrodes |
| $N$ | Number of neurons |
| $J$ | Number of constraints |
| $\Gamma$ | Amplitude response for a single population |
| $w$ | Population weight |
| $d_{\text{norm}}$ | Electrode-population distance |
| $\mathbf{p}'$ | Electrode position |
| $\mathbf{p}$ | Neuron position |
| $\mathbf{P}$ | Population position |
| $v'$ | Voltage at electrode |
| $v$ | Voltage at neuron |
| $V$ | Voltage at population |
| $q'$ | Charge of electrode |
| $q$ | Charge of neuron |
| $Q$ | Charge of population |
| $\mathbf{D}$ | Activity to voltage at electrode transformation matrix |
| $\tilde{\mathbf{D}}$ | Electrode to voltage at population transformation matrix |
| $d$ | Element of $\mathbf{D}$ |
| $\tilde{d}$ | Element of $\tilde{\mathbf{D}}$ |

Our goal in this subsection is to show how the amplitude measured in feedback signals can be related to the synchrony of neural populations. The instantaneous phase Ψ(*t*) and envelope amplitude P(*t*) of a signal *F* (*t*) can be obtained using the analytic signal *R*(*t*)

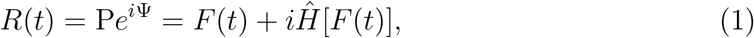

where 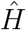 denotes the Hilbert transform. We would like to relate this quantity to those associated with a state of oscillators.

We define the state of *N* regular spiking neurons to be given by the set of oscillators {*θ*_1_(*t*), *θ*_2_(*t*), *θ*_3_(*t*) … *θ_N_* (*t*)}, which are the phases describing where each neuron is in its firing cycle. The phase synchrony of this system can be measured using the order parameter *r*, defined to be

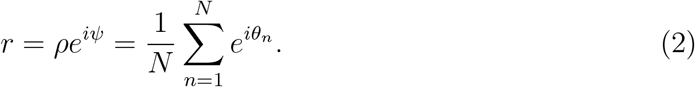

The above definition ensures the magnitude of the order parameter *ρ* can take values between 0 and 1, representing full desynchrony and full synchrony, respectively. We can transform the state of the system to a signal representing the neural activity using a superposition of cosine functions

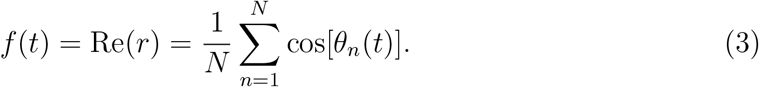

The choice of a cosine function is for mathematical convenience since it corresponds to the real part of (2). In addition to this, the cosine function has a maximum at 0, and in classic coupled oscillator models, phase 0 corresponds to the phase when neurons produce spikes [15]. Hence post-synaptic potentials in down-stream neurons receiving an input from the modelled population will be a smoothed function of spikes produced in phase 0, so the cosine function captures key features of such post-synaptic potentials. Using the Euler relation and comparing (3) with the real part of (2) shows

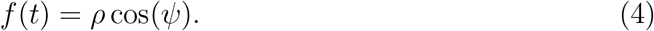

We assume here a simple relationship between the neural activity and feedback signals we may measure, for example tremor and the LFP

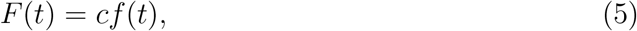

where the experimental signal has now been denoted by *F* (*t*). This is reasonable in the case of ET, where thalamic activity is known to be highly correlated to tremor [5]. Inserting Eq. (5) into (1) gives

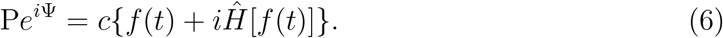

Inserting Eq. (3) into Eq. (6) and using the linearity of 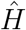 leads to

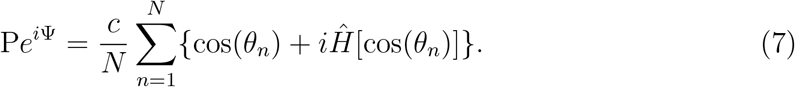

Under the reasonable assumption that the time evolution of *θ_n_* is approximately monotonic, it can be shown that [14]

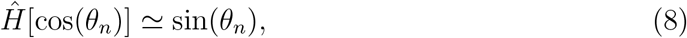

where ‘≃’ is used to indicate ‘approximately equal to’. Therefore

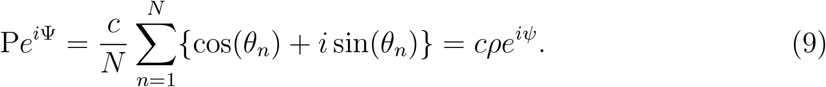

Hence, the instantaneous envelope amplitude and phase (the analytic signal) is relatable to the magnitude and phase of the order parameter using

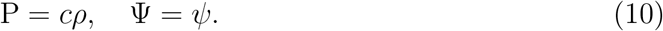

In summary, assuming the experimental data and neural activity are related according to Eq. (5) and that the phases {*θ_n_*} increase monotonically with time, we can use the Hilbert transform of the experimental data to relate the envelope amplitude and instantaneous phase to the magnitude and phase of the order parameter, respectively.

### 2.2. Response Curves

The neuronal phase response curve (nPRC) for a spiking neuron is the change in spike timing due to a perturbation as a function of the inter-spike time. Hansel et al [16] categorised nPRCs into either type I or type II depending on whether a small excitatory (inhibitory) input always advances (delays) a neuron to a next spike or whether it either advances or delays a spike, depending on where the neuron is in its firing cycle, respectively [17]. These effects of inputs can be captured using a simple mathematical function *Z*(*θ*). By mapping where a neuron is in its firing cycle onto a phase variable *θ ∈* [0, 2*π*], the nPRC describes the change in phase of a single neuron due to a stimulus. More precisely, under the assumption of a weak input *ɛU* (*t*), the evolution of a single oscillator can be written in terms of a natural frequency *ω*_0_ in addition to a response term

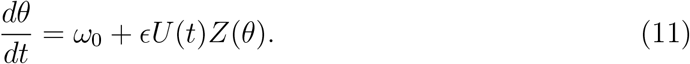

A general neuronal nPRC can be expanded as a Fourier series

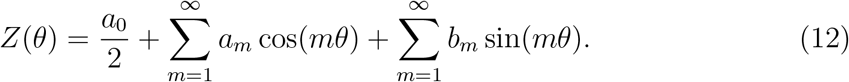

The nPRC type is reflected in the zeroth harmonic *a*_0_, or the shift, with *|a*_0_*|* large and small relative to the other coefficients being indicative of type I and type II curves, respectively. Phase oscillator models which incorporate the nPRC can be shown to reproduce the experimentally-known characteristics of a patient’s response to stimulation [14], namely that the effects should be both amplitude and phase dependent [4, 12]. This leads to the concept of the phase response curve (PRC) and the amplitude response curve (ARC) for feedback signals, such as LFP and tremor, which can be described by perturbing a population of oscillators and respectively describe changes in the phase and amplitude of the feedback signal at the point of stimulation. The instantaneous curves, which are functions of both the phase and amplitude at which the stimulation is delivered, are not commonly found in experimental analysis due primarily to the difficulties associated with obtaining a function of two independent variables from noisy data. It is more common to find the averaged response curves, which are only functions of the phase and are averaged over the amplitude. Such curves are readily obtainable using standard signal processing techniques and have been used to characterise a patient’s response to stimulation [18, 12, 19].

### 2.3. The Kuramoto Model

Modelling the effects of DBS generally poses a challenge since the brain networks involved in disorders such as ET (cortico-thalamic circuit) and PD (cortico-basal-ganglia circuit) are complex and it is still debated from which parts of these circuits the pathological oscillations originate [20, 21]. The task can be made more tractable by considering a simple phenomenological model which does not attempt to explicitly describe the underlying circuits, but rather focuses on general mechanisms leading to the synchronization of neurons. One example of this is the Kuramoto model, [22, 23] where the dynamics of neurons are described using a system of homogeneously coupled oscillators, whose phases evolve according to a set of underlying differential equations. Such models are particularly attractive due to their simplicity and explicit dependence on phase, which makes them convenient for describing the effects of phase-locked stimulation. In the previous section we showed that the oscillation data typically measured in experiment can be modelled using an underlying system of oscillators, whose state is described by the set of *N* phases {*θ_n_*}. We can describe the time evolution of this state (for a single population) using the Kuramoto equations, with an additional term describing the effects of stimulation [22, 7]

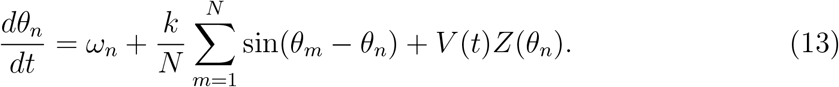

The first term of (13) is the natural frequency *ω_n_* which describes the frequency in the absence of external inputs. The second term describes the coupling between the activity of individual neurons, where *k* is the coupling constant which controls the strength of coupling between each pair of oscillators and hence their tendency to synchronize. The third term describes the effect of stimulation, where the intensity of stimulation is denoted by *V* (*t*). The nPRC denoted by *Z*(*θ_n_*), describes a neuron’s sensitivity to stimulation at a particular phase and reflects the observation that the effects of stimulation depend on where a neuron is in its firing cycle [24]. Using the definition of the order parameter given in Eq. (2), Eq. (13) can be transformed to give

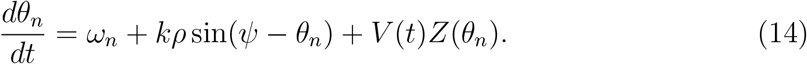

In this form, it is clear that each oscillator has a tendency to move towards the population phase *ψ* and that the strength of this tendency is controlled by the coupling parameter *k*. To gain an intuition for this behaviour readers may wish to explore an online simulation of the model [25].

### 2.4. Reduced Kuramoto Model

In the previous section, we described the dynamics of a finite system of oscillators using the Kuramoto equations given by Eq. (14). In this model, stimulation is described as a perturbation to the phase of an oscillator, with each oscillator experiencing a different effect of stimulation depending on its phase (and determined by *Z*(*θ*)). Stimulation therefore has the effect of changing the distribution of oscillators and hence the order parameter of the system. Since the order parameter, given by Eq. (2), is determined by both the amplitude and phase of the system, the expectation is that stimulation will lead to a change in both these quantities, which we refer to as the instantaneous amplitude and phase response of the system. To obtain analytical expressions for these quantities, we can consider an infinite system of oscillators satisfying the *ansatz* of Ott and Antonsen [26, 27]. In our previous work [14], we showed that for a general nPRC given by Equation (12) and assuming the natural frequencies are Lorentzian distributed with centre *ω*_0_ and width *γ*, the instantaneous change in the order parameter can be written as

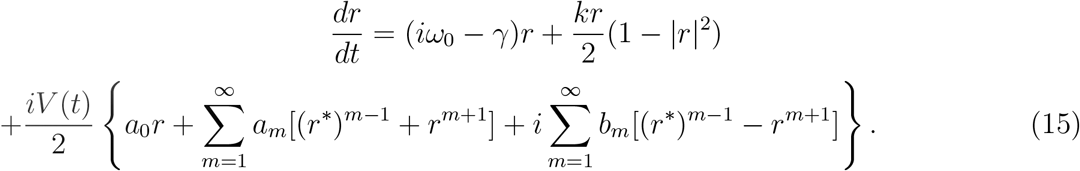

Using this, we can find expressions for the ARC and PRC due to stimulation

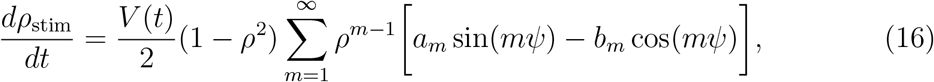

and

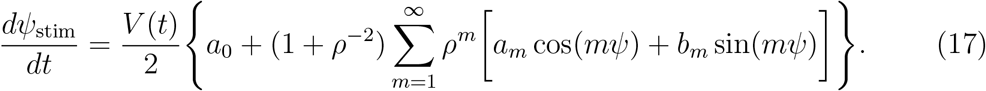

## 3. Reproducing Tremor in ET Patients

We now address the question of whether the Kuramoto model can produce oscillations which are compatible with tremor data from ET patients. To account for random forces which may influence the firing of individual neurons, the Kuramoto model can be extended to include a noise term, which we take here to be a Wiener process. The time evolution for *θ_n_* then becomes

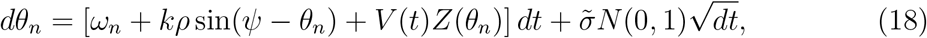

where 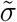 is the noise amplitude and *N* (0, 1) is a random number sampled from a standard normal distribution. During a simulation, the set of oscillators {*θ_n_*} evolves according to (18) and oscillations can be generated using Equation (3). The oscillation data output from the Kuramoto model (18) depends on the choice of parameters {*ω_n_*}, *k*, *N* and 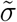. We can characterise these oscillations by using features extracted from the data, which we choose to be: the power spectral density (PSD), the probability density function (PDF) for the amplitude and the PSD of the envelope amplitude. To reproduce the response of a particular patient, we also fit to the averaged PRC [14, 12, 28] of the patient by adjusting the parameters for the nPRC. These characterisations can be applied to both the experimental and synthetic simulated data from the model. The similarity between features from the simulated and experimental data can then be quantified using least squares, which in turn allows us to quantify the degree of similarity between simulated and experimental oscillation data. By using this similarity measure as a cost function, we can then find the parameters of the Kuramoto model which minimise the cost using optimisation and thus find the parameters which allow the model to produce oscillations similar to experimental data.

The computational cost of the optimisation depends on a number of factors, including the number of parameters. To ensure feasibility and prevent overfitting, we choose a reasonable number of oscillators *N* = 60 and sample {*ω_n_*} from a normal distribution with mean 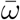 and standard deviation *s_ω_*. We chose to simulate the Kuramoto model with {*ω_n_*} sampled from a normal distribution as opposed to a Lorentzian distribution (which was assumed in the derivation of the response curves) since the long tails of the latter can lead to a non-monotonic evolution for *ψ*, due to sampling small/negative *ω_n_*. This can be problematic for the methodologies used to calculate the phase response curves, which require a monotonic evolution for *ψ*.

We fit the Kuramoto model to tremor data [12, 29] from ET patients deemed to have significant response curves [28]. The parameters found through optimisation are provided in Table 2. Figures 1 and 2(a)-(c) show the Kuramoto model is able to fit well to the features taken from the experimental data. Output from the model can be seen in Figure 3 and shows the resulting simulated data to be quite compatible with that found from experiment. The model can be seen to capture the basic properties of the experimental data, but not the more exotic features, such as the sustained periods of lower amplitudes, which are likely due to non-stationarity. Figures 2(d)-(f) show that the fitted model is able to reproduce the amplitude response for patients generally well, although Figure 2(d) does show a noticeable phase shift between the simulated and experimental curves for Patient 1. Overall, our findings suggest that it is reasonable to use the Kuramoto model as a model for tremor in ET patients. It is on this basis that we derive the expressions for the response curves in subsequent sections. A more detailed description of our fitting methodology, together with details of the experimental data, can be found in the Appendix.

**Figure 1.**
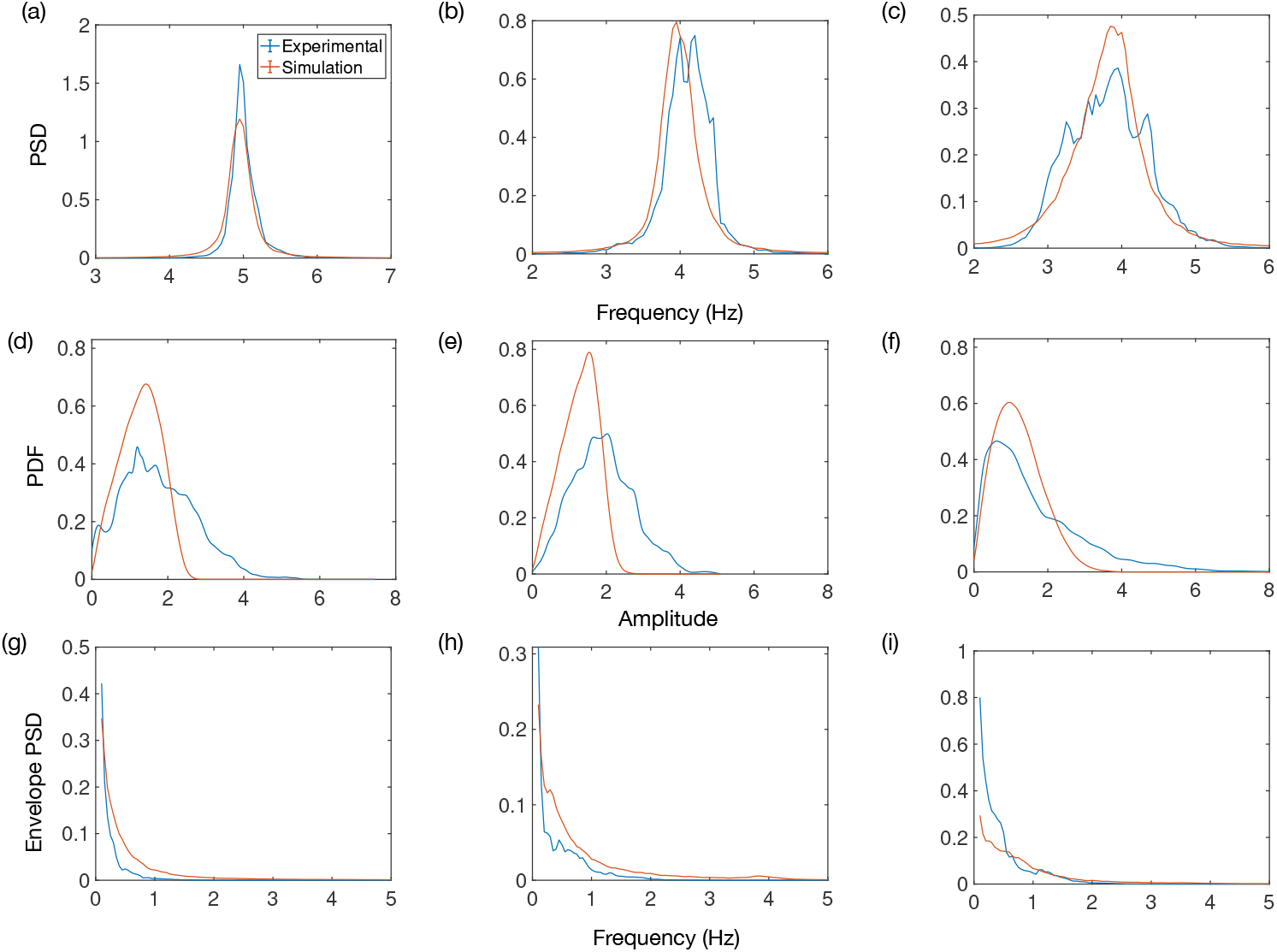
Fits to various features extracted from oscillation data. The first row (a)-(c) is the power spectral density (PSD), the second row (d)-(f) is the probability density function (PDF) for the envelope amplitude and the third row (g)-(i) is the PSD of the envelope. Columns (a)-(g), (b)-(h) and (c)-(i) are for patients 1, 5 and 6, respectively.

**Figure 2.**
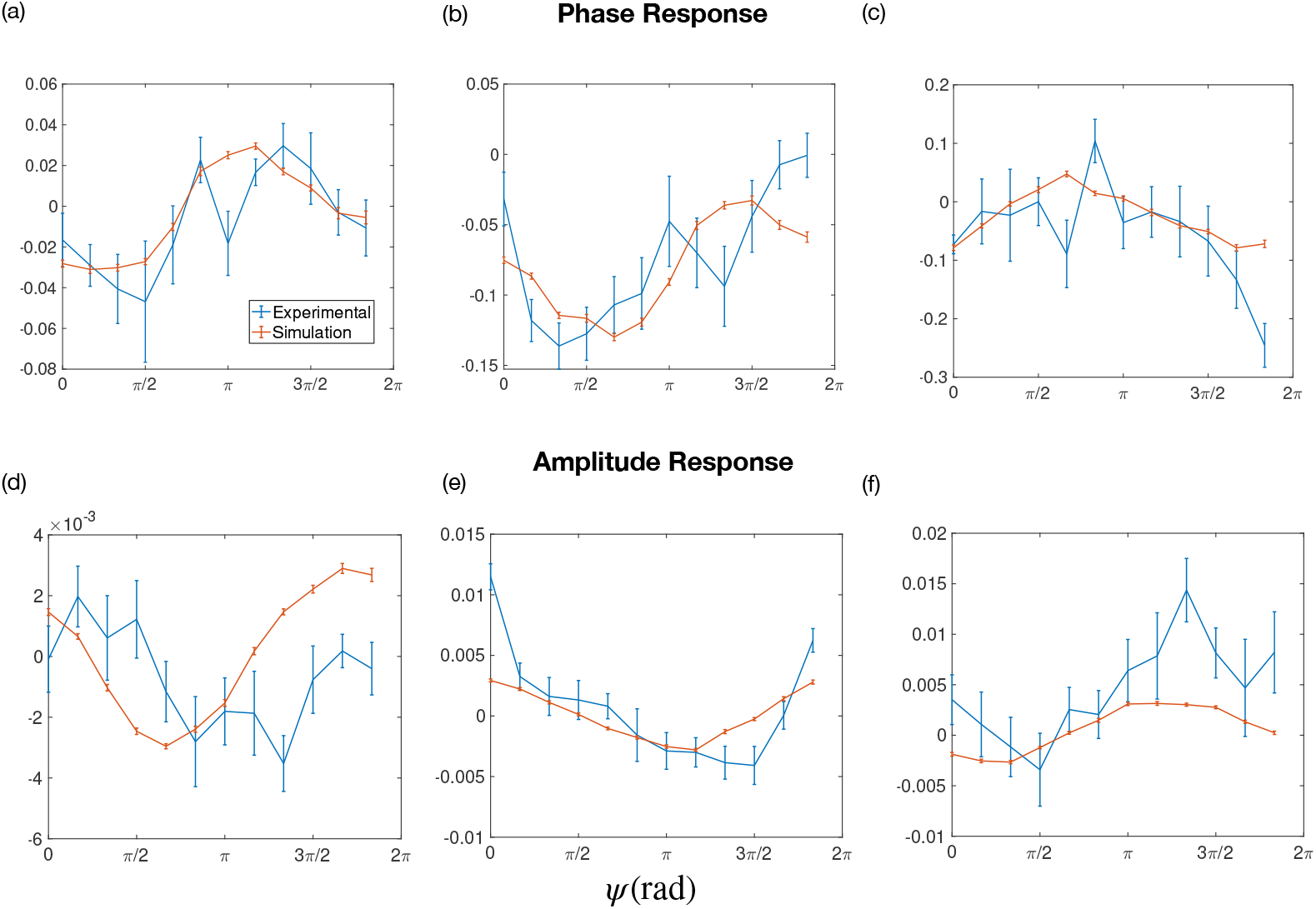
Comparison between the averaged response curves for experimental data and the fitted Kuramoto model. The phase response curve was used as a feature during the fitting procedure. The amplitude response curve is predicted from the model. Columns (a)-(d), (b)-(e) and (c)-(f) are for patients 1, 5 and 6, respectively.

**Figure 3.**
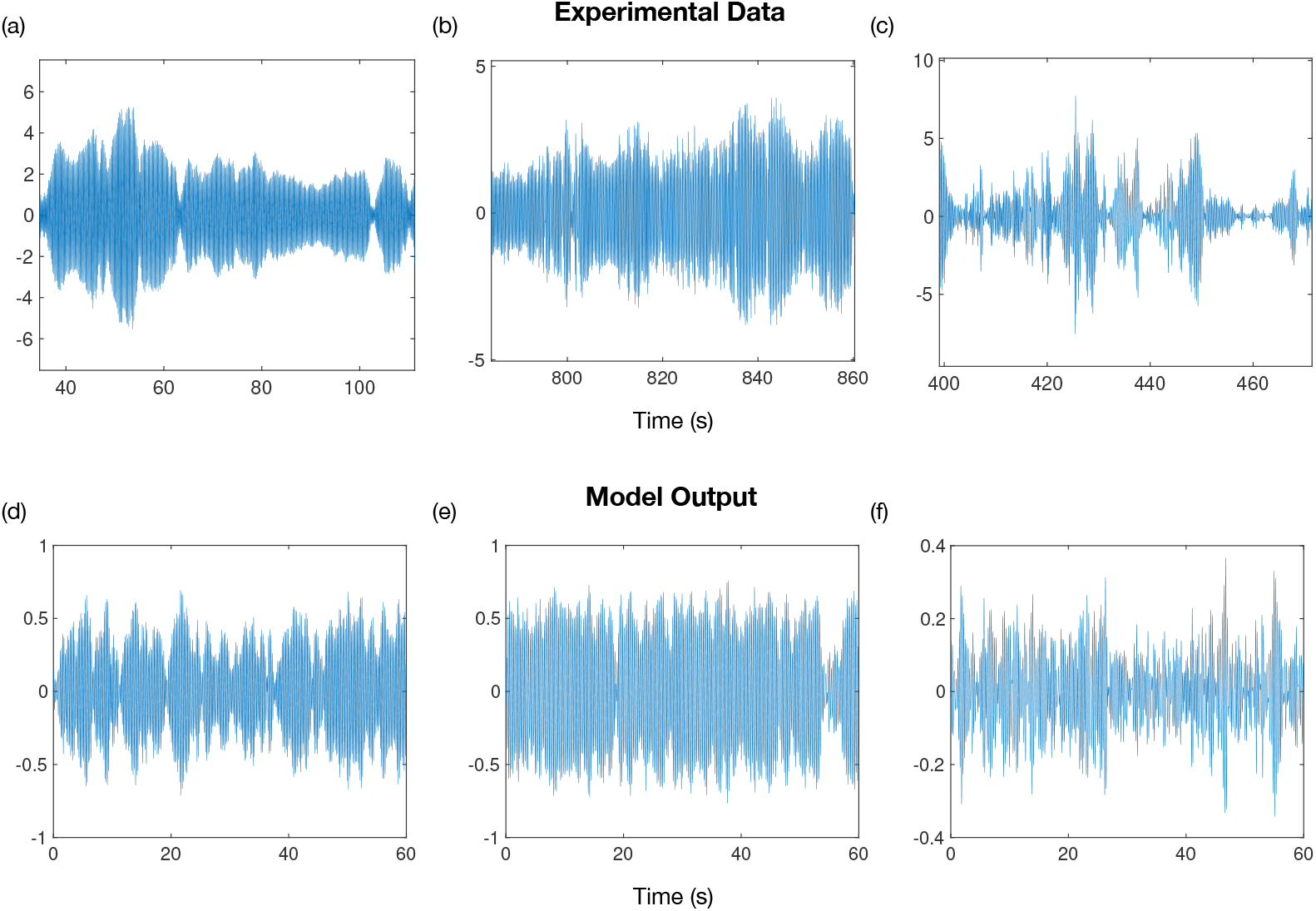
Comparison between experimentally measured tremor data [12] and output from the fitted Kuramoto model. Columns (a)-(d), (b)-(e) and (c)-(f) are for patients 1, 5 and 6, respectively.

**Table 2.**
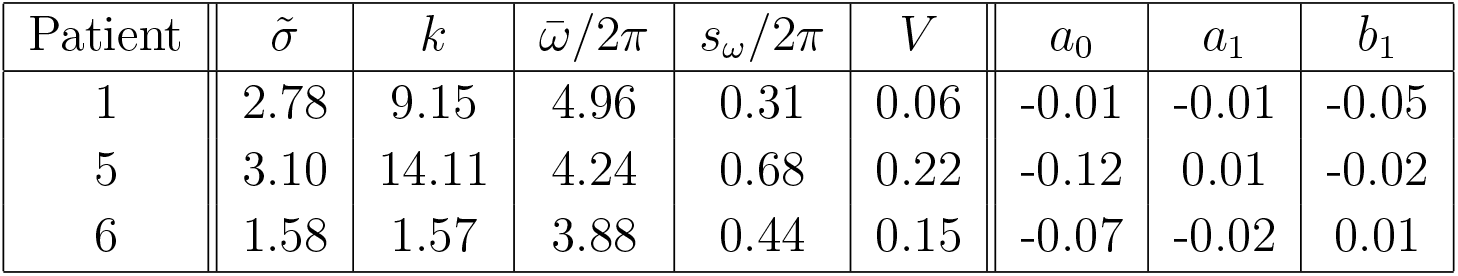
Parameters for the single population Kuramoto model given by Equation (18). The parameters were found by fitting the model to tremor data taken from ET patients by Cagnan et al [12].

| Patient | $\tilde{\sigma}$ | $k$ | $\bar{\omega}/2\pi$ | $s_{\omega}/2\pi$ | $V$ | $a_0$ | $a_1$ | $b_1$ |
| --- | --- | --- | --- | --- | --- | --- | --- | --- |
| 1 | 2.78 | 9.15 | 4.96 | 0.31 | 0.06 | -0.01 | -0.01 | -0.05 |
| 5 | 3.10 | 14.11 | 4.24 | 0.68 | 0.22 | -0.12 | 0.01 | -0.02 |
| 6 | 1.58 | 1.57 | 3.88 | 0.44 | 0.15 | -0.07 | -0.02 | 0.01 |

## 4. Theory of Multi-contact DBS

### 4.1. Multi-population Kuramoto Model

We will show in this section that modelling a symptom due to excessive synchrony of multiple neural populations can be achieved by using a simple extension of the concepts presented in Sections 2.1 and 2.3. The set of oscillators {*θ*_1_(*t*), *θ*_2_(*t*), *θ*_3_(*t*) *… θ_N_* (*t*)} can be arbitrarily divided into *S* populations with *N_σ_* oscillators for the *σ*th population.

The order parameter defined by Equation (2) can then be rewritten using a double summation

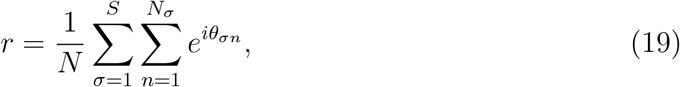

with oscillator *n* of population *σ* being denoted by *θ_σn_*. The factor of 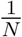 can be brought inside the first summation and rewritten as 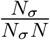. Then, with

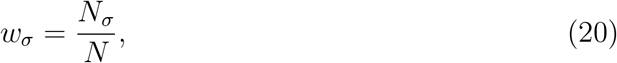

the order parameter for the system can be written as

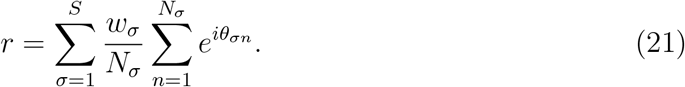

Using the definition of the order parameter (2), Eq. (21) can be written as a weighted superposition of the order parameters for each population

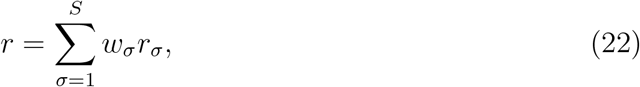

with

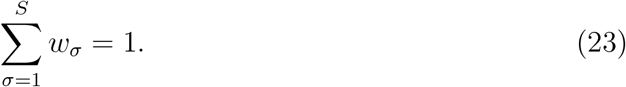

We define *r* to be the global order parameter with amplitude *ρ* and phase *ψ* and *r_σ_* to be the local order parameter for population *σ* with amplitude *ρ_σ_* and phase *ψ_σ_*. The importance of the global order parameter is that its magnitude *ρ* is a measure of total synchrony and hence should be highly correlated to the severity of a symptom, such as tremor in the case of ET. In the case of PD, symptom severity could be measured using the unified Parkinson’s disease rating scale (UPDRS) scores [30]. Therefore, we will consider how to stimulate to maximally reduce the magnitude of the global order parameter.

We can also relate (22) to feedback signals we might measure by using (3) and taking the real part. Under the assumption (5) relating the neural activity to the feedback signal we obtain an expression for the feedback signal in terms of population activities

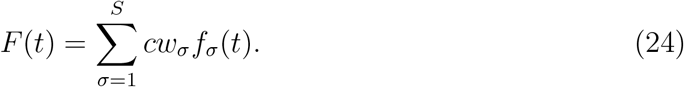

We refer to *F* (*t*) and {*f_σ_*(*t*)} as the global and local signals (or population activities), respectively. Using (4), Equation (24) can also be written in terms of the global and local amplitudes and phases

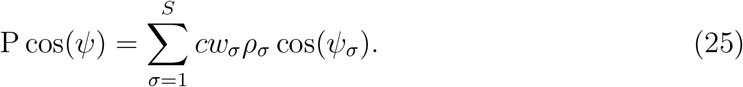

The Kuramoto equations (13) can also be rewritten in terms of the population phases *ψ_σ_* and amplitudes *ρ_σ_*

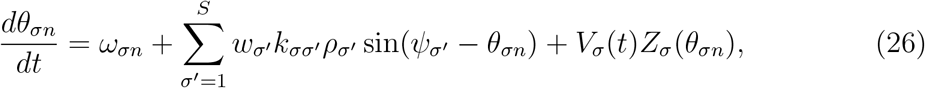

where *V_σ_*(*t*) is the now the stimulation intensity at a population *σ*. The coupling constant *k* in Eq. (13) is now a *S × S* matrix with elements *k_σσ′_*. The diagonal and off-diagonal elements describe the intrapopulation and interpopulation coupling, respectively.

### 4.2. Multi-population Response Curves

We now derive an expression describing the change in the global amplitude due to stimulation as a function of the local (population) amplitudes and phases. For now it is assumed that the local quantities (to base the stimulation on) can be measured. We will discuss how these quantities can be measured later. Using the polar form of the order parameter (2), Equation (22) can be written as a summation involving the amplitudes and phases of individual populations

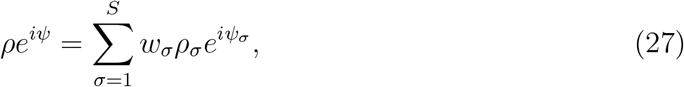

Taking the time derivative of (27) leads to

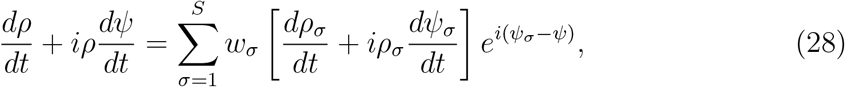

which can be written in terms of the real and imaginary components

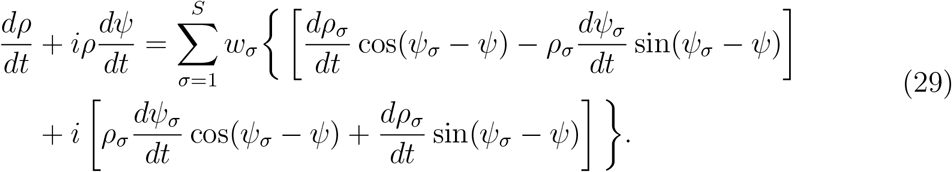

It can be seen that the time derivative of the amplitude is the real part of (29)

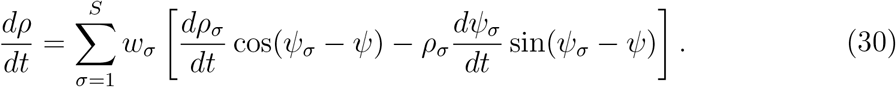

The quantities *dρ_σ_/dt* and *dψ_σ_/dt* of Equation (30) are the changes in the amplitude and phase of a population with respect to time. If we assume the distribution of phases within a population satisfies the *ansatz* of Ott and Antonsen [26], we can substitute Eq. (16) and Eq. (17) into (30) to obtain the amplitude response due to stimulation in terms of the Fourier coefficients of *Z*(*θ*)

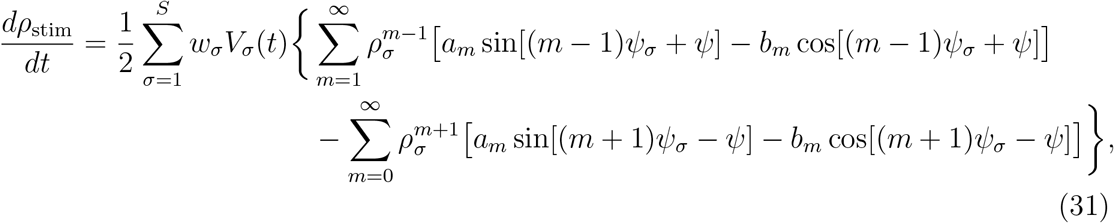

where, for simplicity, we assume that *Z*(*θ*) is the same for all populations. Equation (31) contains an expansion over the harmonics of *Z*(*θ*). In our previous paper, we demonstrated that, for a biologically realistic nPRC, it is reasonable to neglect higher harmonic terms (*m* > 1) [14], leading to a simpler expression for the instantaneous amplitude response

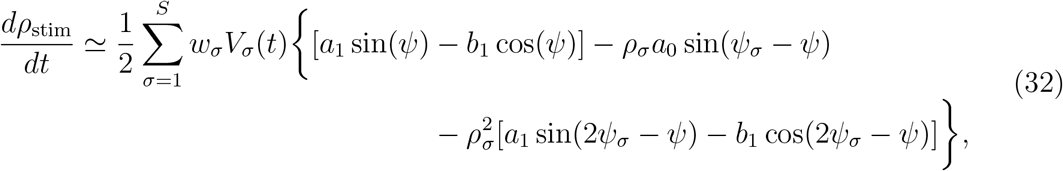

Equation (32) shows the global reduction in amplitude can be expressed as a sum of contributions from each population, with each term dependent on 3 variables: the global phase *ψ*, the local phase *ψ_σ_* and the local amplitude *ρ_σ_*. It also suggests that stimulating on the basis of local quantities may not always be advantageous. It can be seen that the terms of Equation (32) can be divided into two categories: ones which depends on both global and local quantities and ones which depends only on global quantities. The terms depending on both the global and local phases are also dependent on the local amplitudes. In cases where the local amplitude is small, i.e. *ρ_σ_* « 1, we can neglect the term involving 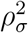, leading to a simplified expression

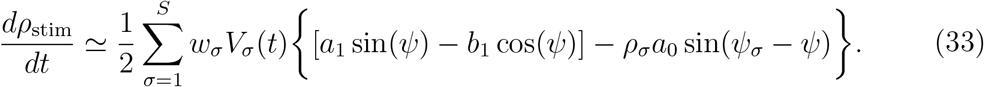

Here, it can be seen that the amplitude response would be dependent only on the global phase if the zeroth harmonic of the nPRC *a*_0_ is negligible, which is the case for type II nPRCs. It can also be seen that the dependency of the amplitude response on the local quantities of population *σ* becomes less at increasingly lower local amplitudes *ρ_σ_*. In addition to this, the dependence on sin(*ψ_σ_ − ψ*) implies that stimulating on the basis of local quantities would only have an effect if the phases of individual populations differ sufficiently from the mean phase. One situation in which such phase difference may be particularly high are for clustered configurations of oscillators. Examples of different configurations of oscillators are shown in Figure 4.

**Figure 4.**
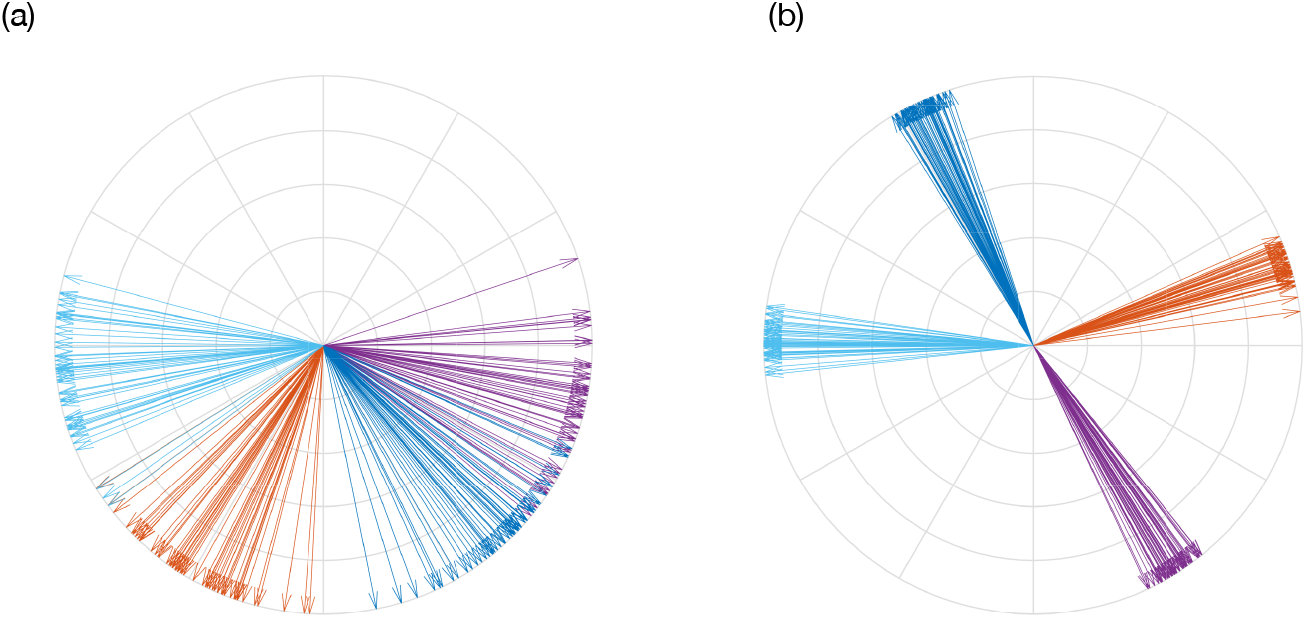
Different configurations of oscillators color coded according to population showing (a) unimodal distribution (b) multimodal (clustered) distribution. Configurations were obtained by simulating the multi-population Kuramoto equations (26).

Plots for the amplitude response (32) together with the corresponding nPRC using the fitted parameters from Table 2 for ET patients 1, 5 and 6 can be seen in Figures 5(a), (b) and (c), respectively. For a given local amplitude, we plot a single term from the summation over populations in Equation (32). This provides the contribution of a single population to the amplitude response as a function of the local and global phases. Regions in blue are areas of amplitude suppression while orange regions predict amplification. In both cases, these regions can be seen to occur in bands. Graphically, the dependence of the amplitude response on the global and local phases can be inferred from the direction of the banding. A purely horizontal band implies the amplitude response is independent of the local phase. An example of this can be seen at low amplitudes in Figure 5 (a). Other plots show diagonal banding, which implies the amplitude response is dependent on both the global and local phases. This behaviour can be understood by considering the 3 terms of (32). At low amplitudes, the first term dominates, which is only dependent of the global phase. As the local amplitude increases, the second and third terms depending on local quantities become increasingly more important. For the cases where *|a*_0_*|* is small, the effect is less apparent. The left panel of Figure 5 (a) shows that stimulation can either increase or reduce the phase (i.e. an nPRC of type II), implying a relatively small *|a*_0_*|*. Hence, for this patient, the second and third terms are negligible, except at higher amplitudes. Figures 5 (b) and (c) shows that stimulation has the effect of only increasing the phase, which is indicative of *Z*(*θ*) with larger *|a*_0_*|*. For these systems the amplitude response can be seen to depend more strongly on the local phase for all amplitudes.

**Figure 5.**
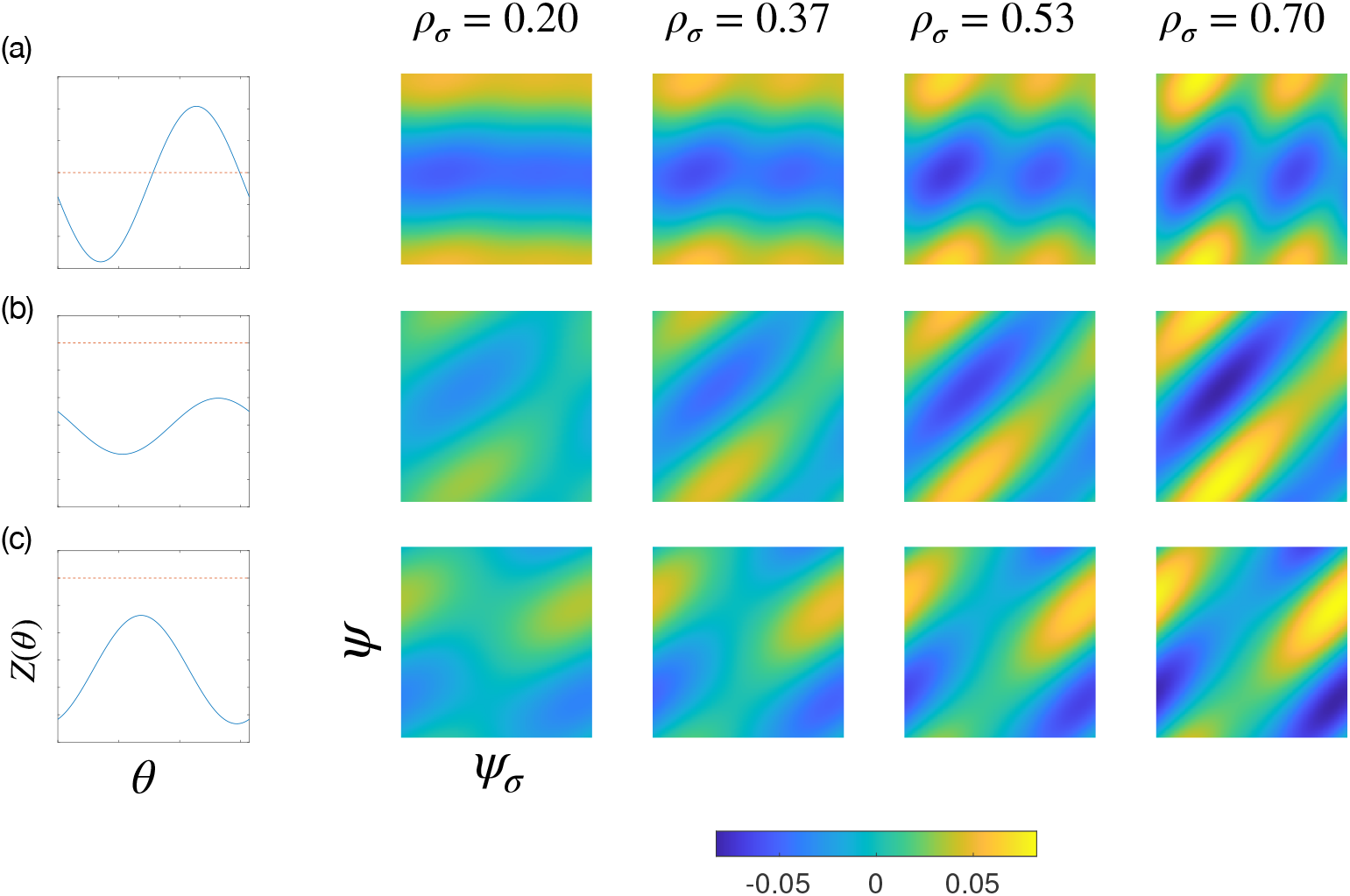
The predicted contribution of a single population to the amplitude response at different local amplitudes *ρ_σ_* according to Eq. (32). Each panel corresponds to a single ET patient from the study of Cagnan et al [12], where the Fourier coefficients of the nPRC were determined using a fitting procedure. Panels (a) (b) and (c) are for patients 1, 5 and 6, respectively. For each plot, the vertical axis is the global phase (*ψ*) and the horizontal axis is the local (or population) phase (*ψ_σ_*). The corresponding nPRC *Z*(*θ*) is also shown, with zero indicated by a red dashed line. Blue regions indicate areas where stimulation is predicted to suppress amplitude.

### 4.3. Obtaining Population Activities Through Electrode Measurements

In this subsection, we will describe how the local phases {*ψ_σ_*} and amplitudes {*ρ_σ_*} can be recovered using LFP measurements through different contacts. This requires us to incorporate information about the geometry of the electrode placement into the equations for the response curve in addition to assigning a physical interpretation to the population activity. Our aim here is not to construct a detailed electrophysiological model of neuronal activity but rather to present a very general form for the voltage measured at an electrode contact. We formulate our expressions here in terms of electric charge, but the same form also permits the use of currents. In addition to this, our expressions include summations over neurons, but an equally valid expression can be made by summing over elements of space, as is the case in multi-compartmental models [31]. The quantities we consider in our model are voltages 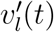 measured at electrode *l* due to the activity of population *σ* producing charges *Q_σ_*(*t*) and voltages *V_σ_*(*t*) at population *σ* due to stimulation which delivers charge 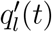 to electrode *l*. The voltage *V_σ_*(*t*) can also be thought of as the ‘stimulation intensity’ experienced at population *σ*.

We begin by considering a system of *L* electrodes and *N* neurons with positions in space denoted by **p**′ and **p**, respectively. From now on, we will use the following notation throughout: primes to denote quantities associated with electrodes, lower case for neuronal quantities and upper case for population quantities. Voltages measured at an electrode arise due to the geometry of the electrode-neuron system and the intrinsic electrical activity of each neuron. We express the voltage measured at an electrode in terms of a summation over charges due to the neurons *q_n_*(*t*)

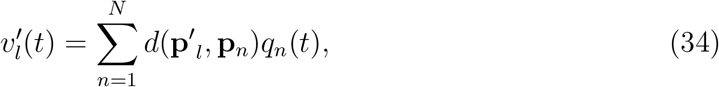

where 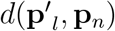 are coefficients which reflect the medium and geometry of the electrode-neuron system. For example, in the case of a coulombic system, the coefficients would be

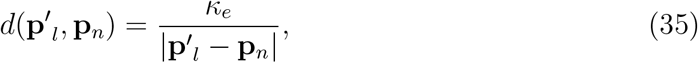

where *κ_e_* is the Coulomb constant. As before, a system of neurons can be arbitrarily divided into *S* populations, with each neuron referenced by both a population and position index *σ* and *n*, respectively.

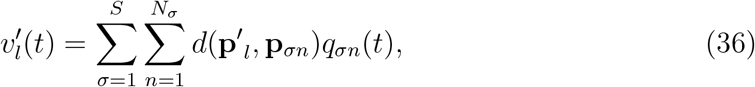

We now let **p**_*σn*_ = **P**_*σ*_ + Δ**p**_*σn*_, i.e. we now define a vector to a neuron in terms of a vector to a region (or population) plus a shift.

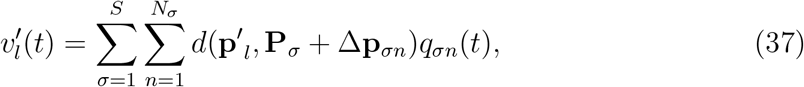

If we assume the region at **P**_*σ*_ to be small, then

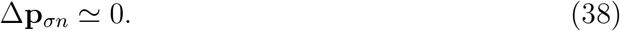

The potential at the electrode can then be written in terms of population activity

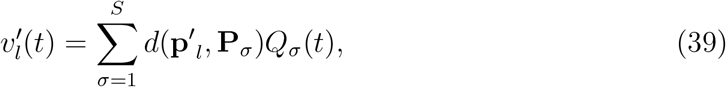

where

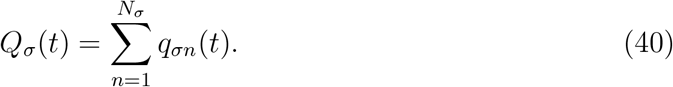

The time dependent charge of a population *Q_σ_*(*t*) can be related to the neural activity by assuming a form for *q_σn_*(*t*), specifically that

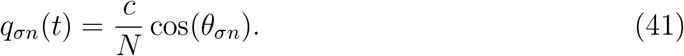

Inserting this into (40) and using (20) gives

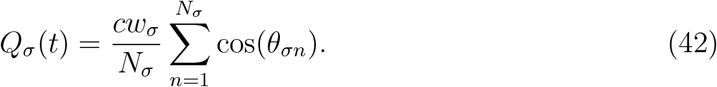

Using (3) and (4) gives an expression for the time dependent charge of a population in terms of the population phase and amplitude

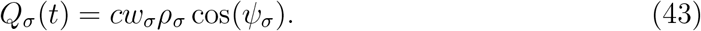

Using (39) the potential at the electrodes can therefore be written in matrix form

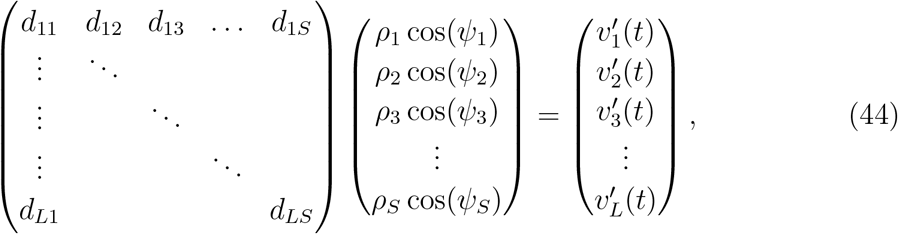

where for simplicity we have denoted 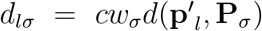. Equation (44) can be expressed in a more compact form with **D** denoting the matrix of coefficients (of dimensions *L × S*), **f** as the vector of neural activities and **v**′ as the vector of electrode measurements.

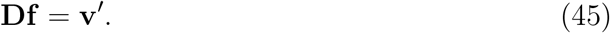

Equation (45) relates the voltages at the electrodes **v**′ to the neural activities **f**. In general, our ability to use Equation (32) in a closed-loop DBS strategy depends on being able to accurately measure the population quantities {*ρ_σ_*} and {*ψ_σ_*}. Equation (44) shows that what we actually measure at the electrodes is a linear superposition of population activities. For the cases where **D** is approximately diagonal, the population quantities could be accurately recovered (although *ρ_σ_* would be scaled). Such cases would represent systems consisting of small separated regions of activity, with each electrode positioned close to each region (see Figure 6(a)).

**Figure 6.**
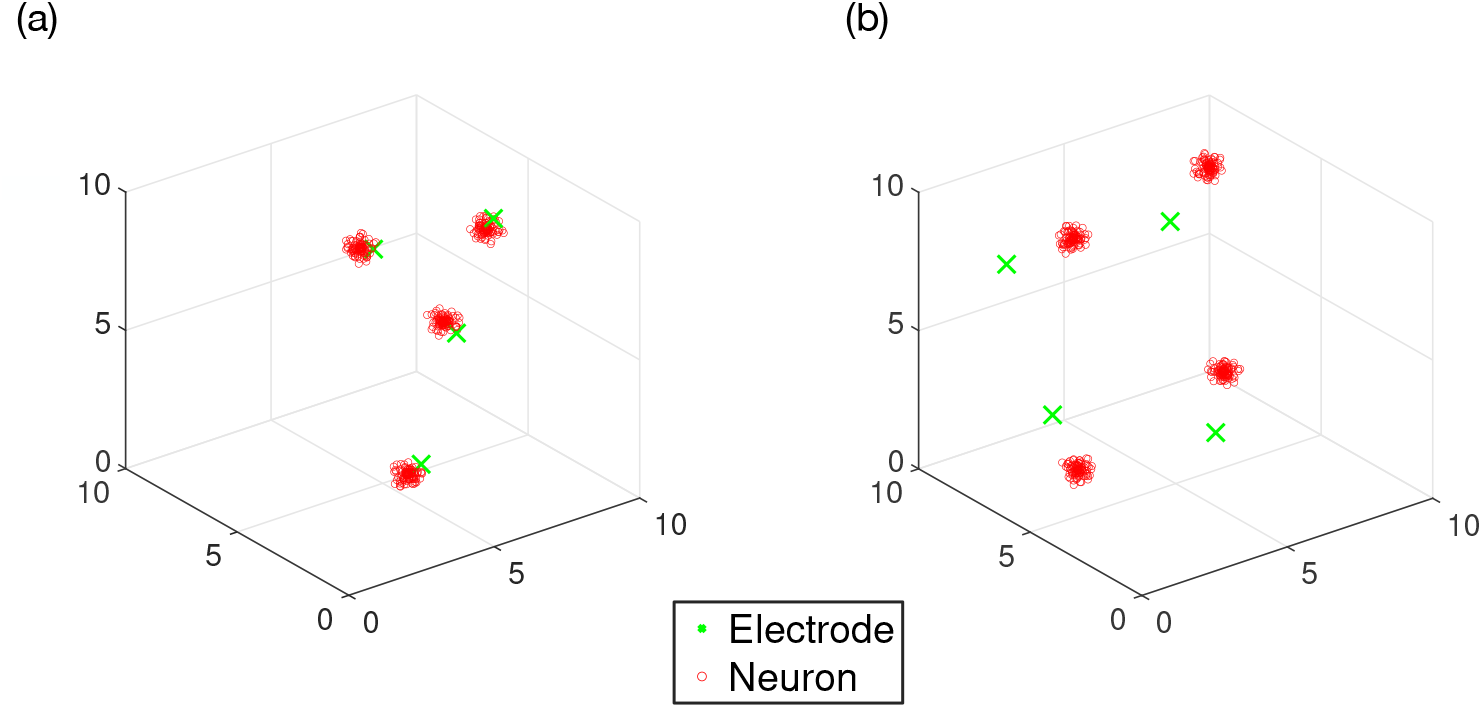
Visualisations of 4 electrode 4 population systems, where each population occupies a small spatial region. Each system was generated by randomly choosing the coordinates of the 4 populations so that they lie within a box of length *L*_box_ = 10. Each electrode is then placed *d*_norm_ distance from a population. Panel (a) shows a configuration where each electrode is placed very close to a population (*d*_norm_ = 0.5). Panel (b) shows a different system (*d*_norm_ = 2) where both the electrodes and populations are more ‘dispersed’. In this scenario, electrodes may record activity from multiple populations.

Methods such as independent component analysis (ICA) [32] are well-suited to solving the general problem of recovering a vector of ‘source signals’ **f**(*t*) (in this case the population activities) given a vector of recordings **v**′(*t*), as expressed in Equation (44), although the method cannot recover the scaling. We consider the special case of a single contact recording, i.e. with *L* = 1, in the appendix. Since in theory the matrix *D* should not evolve with time, we envisage ICA being applied offline to recover *D* and then used to obtain the local signals. In practice, after determining the local signals, Equation (25) should be used to construct the global signal. In this process, the weights {*w_σ_*} should be chosen to give a global signal with an amplitude that is highly correlated to the symptom severity.

### 4.4. Optimal Stimulation Strategy

The equations for the amplitude response (31), (32) and (33) depend on the stimulation intensity at a population *V_σ_*. It is implied, therefore, that the ‘population’ exists at some region in space and that *V_σ_* should take into account the geometry of the electrode placement, how electric fields behave within brain tissue and the charges on a particular electrode. In this subsection, our aim is to incorporate these ideas into an expression for the amplitude response.

Equations (31), (32) and (33) all involve summations over populations, with each term being the product of a weight *w_σ_*, a stimulation intensity *V_σ_* and some intrinsic response, which we shall denote here by Γ_*σ*_. For example, in the case of Equation (31) Γ_*σ*_ would be

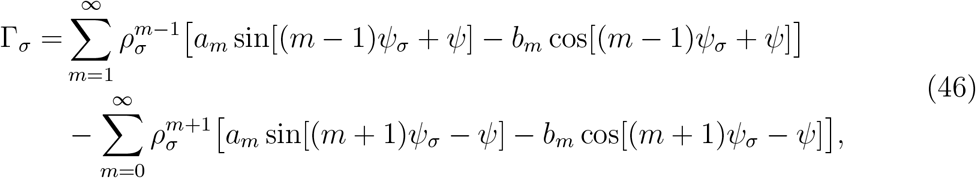

Using this, a more compact expression for the amplitude response can be written using linear algebra notation, with **Γ** equal to the vector of responses and **V** equal to the vector of voltages at a population

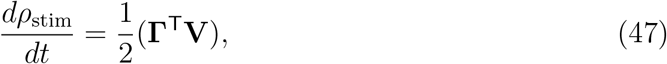

where the weights *w_σ_* are now considered as part of the response **Γ**. The amplitude response involves a ‘stimulation intensity’ **V**(*t*)- an abstract quantity which, intuitively, should not only depend on the charge characteristics at the electrode, but also the geometry of the electrode placement and the properties of the brain tissue. Taken altogether, the stimulation intensity is better interpreted as the voltage at a population, which can be expressed as a weighted superposition of charges at the electrodes

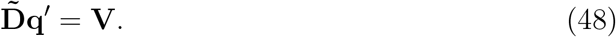

As before, the elements of matrix 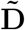 (of dimensions *S × L*) are coefficients which reflect the medium and geometry of the electrode-neuron system. Its worth noting here that Equations (45) and (48) can also be used to model systems where the stimulating and recording electrodes are different, since 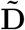 is allowed to be different from **D**^T^. Inserting (48) into (47) leads to an expression for the amplitude response in terms of the charges at the electrodes, i.e. the control variables

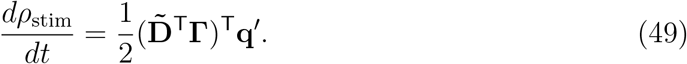

The quantity 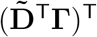 is defined for each time step so that the optimisation becomes a problem of choosing **q**′ so as to minimise *dρ*_stim_*/dt*. Often, concern for tissue damage due to stimulation imposes a limit on how much charge can be delivered to a single or group of contact(s). To account for this and ensure feasibility, we impose two constraints. The first constraint ensures the charge for a particular contact does not exceed some maximum value 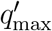

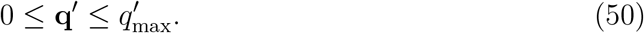

A simple optimal solution (per time step) for Equations (49) and (50) can be found by setting the charge for the *l*th contact to 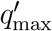 if the *l*th component of 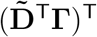 is negative. A second constraint ensures the charge density within a region does not become dangerously high

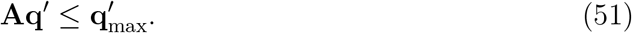

Here, for *J* groups of contacts, the constraint matrix **A** has dimension *J × L* and can be used to constrain the collective charges of the group. The *J*-dimensional vector 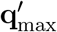 specifies the maximum charge for a particular group of contacts. Equations (49), (50) and (51) are in the standard form for a linear program and are solvable in polynomial time.

## 5. Numerical Simulations

The instantaneous response tells us how the amplitude of a system should change as a function of its state variables but did not take into account the dynamics of the system, such as the coupling which acts to resynchronise the oscillators and the effects of a finite number of oscillators– the latter leading to a breakdown in the underlying assumptions which lead to Equation (32). To better assess the real world performance of a particular stimulation strategy we use the time-averaged response, which requires us to simulate a system using equations (4), (27) and (26).

### 5.1. Simulated systems

We define a system in terms of its electrode-population configuration, dynamics and intrinsic response to stimulation *Z*(*θ*). To construct a particular system we first randomly choose the coordinates of *S* populations such that they lie within a box of length *L*_box_ = 10. We then assign to each population an electrode, which we place *d*_norm_ distance from the population. For a sufficiently large *L*_box_, *d*_norm_ can be used to characterise the system– a small *d*_norm_ means the effects of stimulation are localised to a particular population and increasing *d*_norm_ increasingly delocalises the effects of stimulation. For simplicity, we consider a system consisting of *S* = 4 populations and *L* = 4 electrodes. The analytical expressions for the response curves are for an infinite system, so we require that *N_σ_* is large. For each population, we choose the number of oscillators *N_σ_* = 200 to satisfy this and to remain computationally feasible. We also assume a coulombic system, where each electrode is able to simultaneously record and stimulate. In this case, the elements of *D* are given by

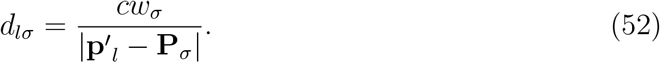

We denote the elements of matrix 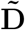 as 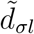, which can be related to **D** using the transpose

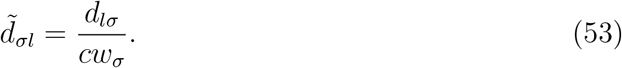

The dynamics of a system are determined by the parameters of the multipopulation Kuramoto model with an additional noise term

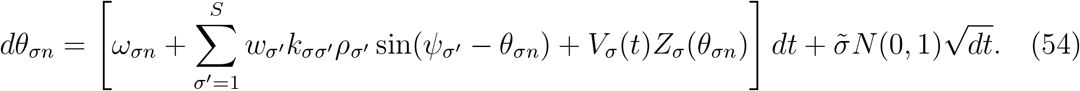

To simplify our testing, we fix the basic parameters of (54) to those found from fitting to Patient 5. As previously mentioned, the natural frequencies {*ω_σn_*} are sampled from a normal distribution. Such simulations represent a greater test for the robustness of the predicted amplitude response due to stimulation (32), which assumes a Lorentzian distribution for {*ω_σn_*}.

The *S × S* coupling constant matrix can be simplified by focussing only on the diagonal and off-diagonal components, which we denote by *k*_diag_ and *k*_offdiag_, respectively. We fix *k*_offdiag_ = 6, so that *k*_diag_ can be used to control the level of clustering for a particular configuration of oscillators-increasing *k*_diag_ leading to increasingly multi-modal distributions of oscillators. The nPRC *Z*(*θ*) was also chosen according to parameters fitted to Patient 5, but we allow the zeroth harmonic *a*_0_ to vary.

### 5.2. Running the simulation

To test each strategy we first create a system according to the set of parameters {*d*_norm_,*k*_diag_,*a*_0_} then choose a stimulation strategy from CR, phase-locked (PL) and ACR. Our implementation of PL stimulation is to use Equation (33), but to neglect all the local terms, which is equivalent to setting *ρ_σ_* = 0.

We use the time-shifted variant of CR neuromodulation [7, 33] in our testing. For a given electrode, stimulation is delivered in bursts of HF pulse trains. The stimulation pattern is time-shifted across each electrode indexed by *l* by

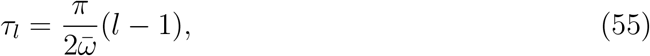

where 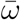 is the mean of the natural frequencies (≃ 4.2 Hz). The number of bursts per second, the burst frequency *f*_burst_, was chosen to be equal to 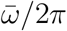 and the HF pulse train frequency *f*_train_ was chosen to be 130 Hz. The width of each burst *t*_burst_ was chosen to be 0.1 seconds. Tass et al originally tested CR on a homogeneously coupled system with *s_ω_* = 0 [7]. We test our implementation and reproduce these results by constructing a simple homogeneously coupled system according to the parameters of Patient 5 given in Table 2, but with 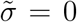, *s_ω_* = 0 and the parameters of *Z*(*θ*) scaled by a factor of 10. The simulation parameters were chosen according to Table 3. The desynchronising effects of CR neuromodulation on this system are shown in Figure 7, which reproduces the results of Tass et al [7].

**Figure 7.**
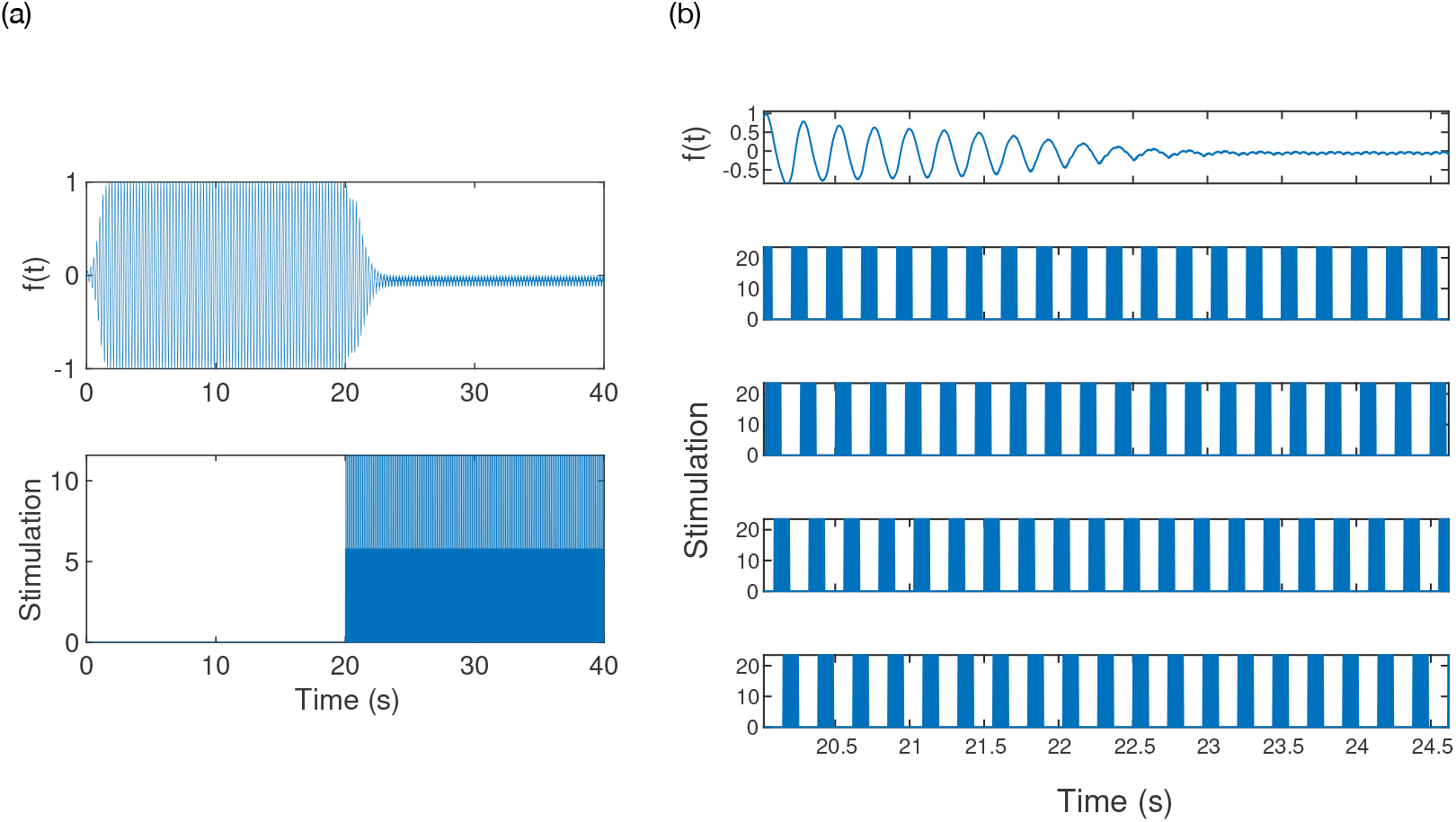
Output from numerical simulations showing the effects of Coordinated Reset (CR). Stimulation was turned on at *t* = 20 seconds. The top panel of (a) shows the model output for a system simulated according to Equations (54) and (24). The bottom panel of (a) shows the stimulation delivered as a function of time, taken to be the average of the charges across the contacts. The bottom panel of (b) shows the stimulation across each contact, with the corresponding model output provided in the top panel.

**Table 3.** Summary of fixed parameters used in the simulations.

| Parameter | Value | Description |
| --- | --- | --- |
| $\Delta t$ | 0.04 | Integration time step |
| $T$ | 2 | Simulation time |
| $L_{\text{box}}$ | 10 | Box length |
| $L$ | 4 | Number of electrodes |
| $S$ | 4 | Number of populations |
| $N_{\sigma}$ | 200 | Number of oscillators per population |
| $k_{\text{offdiag}}$ | 6 | Off-diagonal of coupling constant matrix |
| $d\theta_{\text{max}}$ | $0.2\pi$ | Maximum angle moved per stimulation pulse |
| $n_{\text{trials}}$ | 24 | Number of trials |
| $f_{\text{max}}$ | 130 | Maximum frequency for PL stimulation |
| $f_{\text{burst}}$ | 4.2 | CR burst frequency |
| $f_{\text{train}}$ | 130 | CR HF train frequency |
| $t_{\text{burst}}$ | 0.1 | CR burst time |
| $\bar{\omega}/2\pi$ | 4.2 | Mean oscillator frequency |
| $s_{\omega}/2\pi$ | 0.7 | Standard deviation of oscillator frequency |

The maximum charge for an electrode 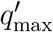 is chosen so that, for a given system, the maximum perturbation to a single oscillator is *dθ*_max_ = 0.2*π* rad. For our testing, we do not constrain ACR using Equation (51), which leads to trivial optimal solutions to the linear program (49) and (50), where the charge for the *l*th contact is set to 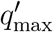 if the *l*th component of 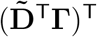 is negative. ACR was tested at 3 maximum stimulation frequencies: 5 Hz, 50 Hz and 130 Hz. The maximum stimulation frequency for PL was fixed at 130 Hz. Equation (54) was then integrated using Euler’s method with a time step of Δ*t* = 0.04 seconds and simulated for *T* = 2 seconds. The PL and ACR strategies were applied according to phases and amplitudes obtained directly from the simulation.

During a simulation, a stimulation pulse is calculated as the average of the charges *q′*(*t*) across the *L* electrodes. Two quantities are calculated after each simulation: the time-averaged value of *ρ*, 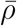 and the total of all stimulation pulses *E* delivered. The former is indicative of the efficacy of the strategy while the latter is related to the total energy consumption of a strategy, which can be used to gauge efficiency. For each set of parameters, the simulations are repeated over 24 trials, with a new electrode-population configuration being generated according to *d*_norm_ for each trial. The parameters *d*_norm_ and *k*_diag_ were chosen within the range *d*_norm_ *∈* [0.1, 6] and *k*_diag_ *∈* [5, 150]. Example output from these simulations, showing the effects of applying ACR, is provided in Figure 8. When compared to Figure 7, it is clear that the stimulation pattern from ACR is significantly different from that produced by CR, with the latter pattern being simply time-shifted across electrodes. The stimulation pattern from ACR allows for the possibility that multiple electrodes may be stimulated simultaneously. A summary of the parameters used in these simulations is provided in Tables 3 and 4.

**Figure 8.**
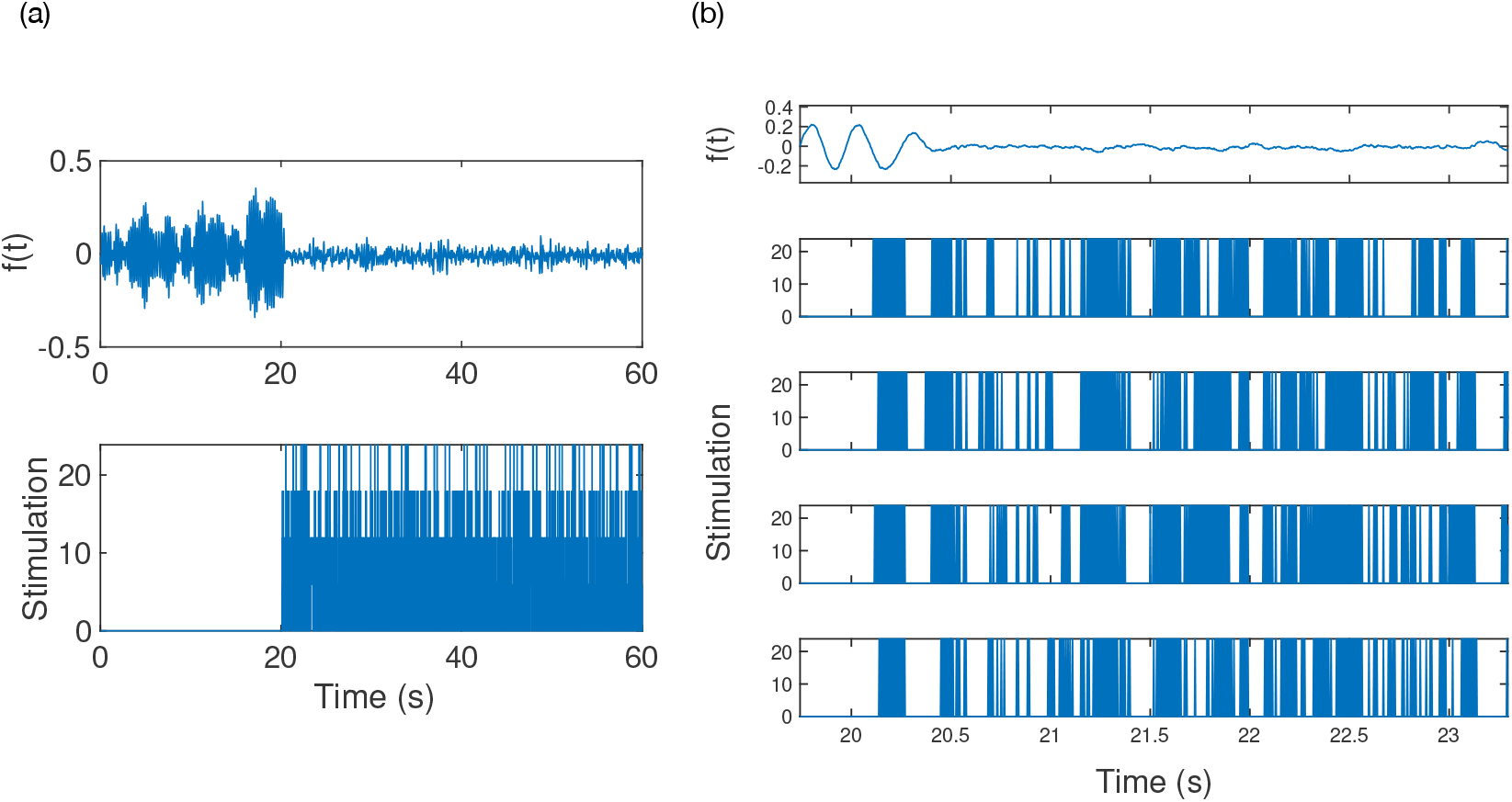
Output from numerical simulations showing the effects of Adaptive Coordinated Reset (ACR). Stimulation was turned on at *t* = 20 seconds. The top panel of (a) shows the model output for a system simulated according to Equations (54) and (24). The bottom panel of (a) shows the stimulation delivered as a function of time, taken to be the average of the charges across the contacts. The bottom panel of (b) shows the stimulation across each contact, with the corresponding model output provided in the top panel.

**Table 4.** Summary of variable parameters used in the simulations. Each parameter was chosen in the range [Min,Max] using a uniform grid of spacing (Max-Min)/*N*_grid_.

| Parameter | Min | Max | $N_{\text{grid}}$ | Description |
| --- | --- | --- | --- | --- |
| $d_{\text{norm}}$ | 0.1 | 6 | 5 | Distance from population to electrode |
| $k_{\text{diag}}$ | 5 | 150 | 5 | Diagonal of coupling constant matrix |
| $a_0$ | 0 | 1.7 | 4 | Zeroth harmonic of $Z(\theta)$ |

### 5.3. Results

Figure 9 shows plots for the average amplitude 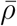, i.e. *ρ* averaged over all trials and all values of *d*_norm_ for a particular value of *k*_diag_ and zeroth harmonic *a*_0_. ACR was tested at maximum frequencies of 130 Hz, 50 Hz and 5 Hz. The maximum frequency of PL was fixed at 130 Hz. The rise in 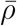 shown between *k*_diag_ = 50 and *k*_diag_ = 70 is indicative of a bifurcation, which is typical in Kuramoto systems [26]. Significant improvements with ACR over PL and CR are observed in simulations when stimulation is delivered at higher frequencies and when *a*_0_ is non-negligible. The utility of ACR over other methods is also shown to be greatest when *k*_diag_ is larger, which corresponds to larger local amplitudes *ρ_σ_* and increased clustering.

**Figure 9.**
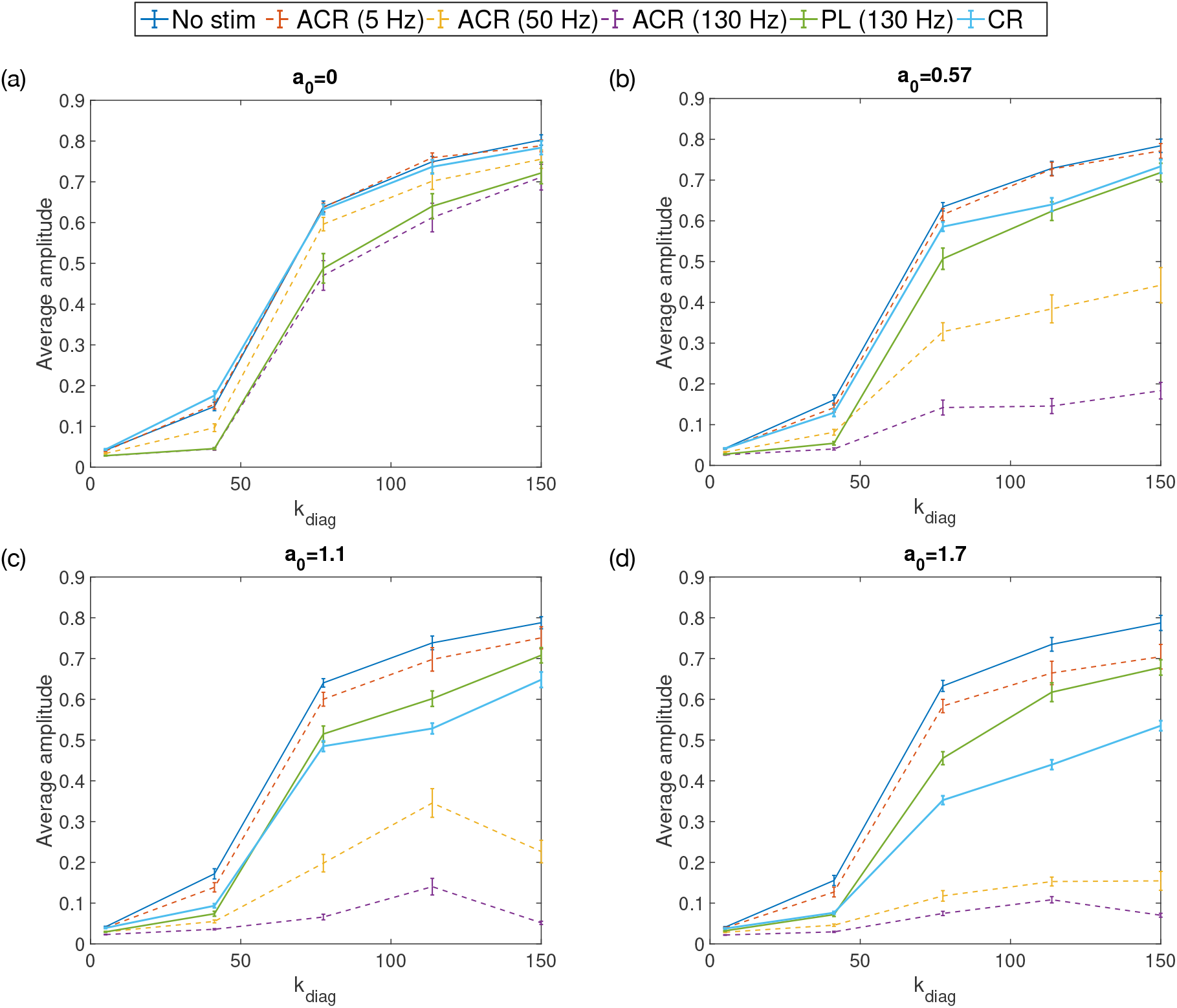
The average amplitude of a simulated Kuramoto system with a coupling constant *k*_diag_ for different stimulation strategies: no stimulation (no stim), Adaptive Coordinated Reset (ACR), phase-locked (PL) and Coordinated Reset (CR). The maximum stimulation frequency used for ACR and PL is also given in the legend. Dashed lines are for the ACR method. Each sub plot shows a set of simulations performed with a particular zeroth harmonic of the nPRC *a*_0_.

The efficacy of CR can also be seen to improve with systems with larger *a*_0_. With *a*_0_ = 0, CR and no stimulation are shown to be equally effective. The results shown in Figure 9 are in contrast with those shown in Figure 7, indicating that the efficacy of CR is sensitive to the parameters 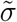 and *s_ω_* in addition to the scaling of *Z*(*θ*).

For systems where *a*_0_ ≃ 0, PL and ACR are found to be equally as effective, as predicted. The efficacy of ACR at 130 Hz is greater than at other frequencies, but with more energy usage than PL and similar energy usage to CR, as shown in Figure 10. ACR at 50 Hz is found to have good efficacy for *a*_0_ > 0 but with significantly less energy usage than PL and CR. ACR with low frequency stimulation at 5 Hz (or approximately the tremor frequency) is predicted to have little to no effect on all the systems tested.

**Figure 10.**
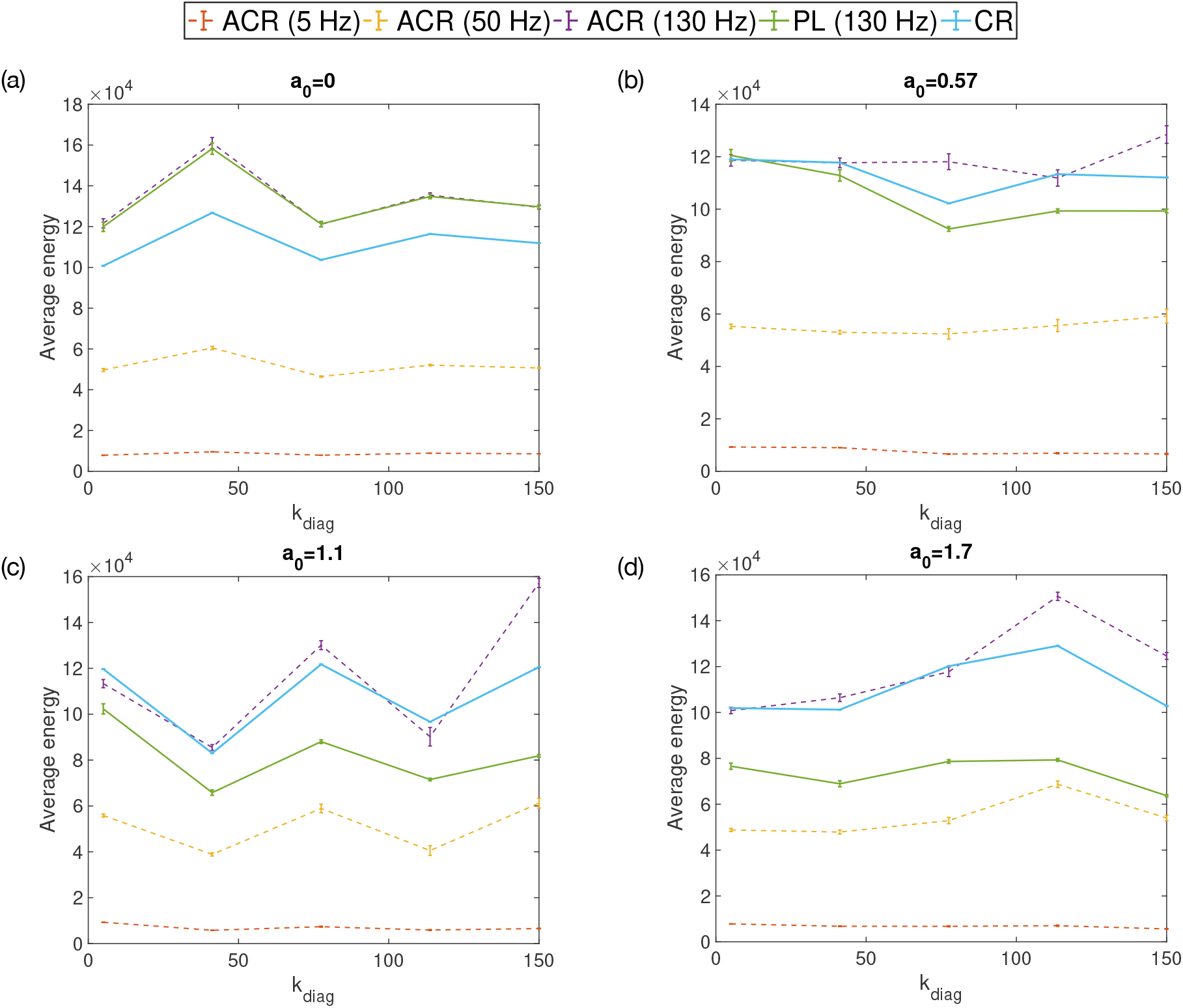
The average energy used by: no stimulation (no stim), Adaptive Coordinated Reset (ACR), phase-locked (PL) and Coordinated Reset (CR) stimulation strategies on a simulated Kuramoto system with a coupling constant *k*_diag_. The maximum stimulation frequency used for ACR and PL is also given in the legend. Dashed lines are for the ACR method. Each sub plot shows a set of simulations performed with a particular zeroth harmonic of the nPRC *a*_0_.

As previously mentioned, we expect the utility of ACR to be greatest for those systems described by type I nPRCs as the amplitude response curve then depends explicitly on non-negligible terms involving population quantities. Equation (32) shows that when *a*_0_ = 0, the terms involving population quantities depend on second harmonics 2*ψ_σ_*. These terms carry a factor of 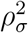 and hence are likely to be negligible, except when *ρ_σ_* is reasonably large. To investigate these effects, we simulate systems with *a*_0_ = 0 and use a larger stimulation intensity 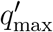. By comparing ACR to PL, we can then ascertain the impact of these second harmonic terms. A comparison of the efficacy of ACR with PL and CR, is shown in Figure 11 for different stimulation amplitudes 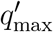 and with *a*_0_ = 0. Increasing stimulation amplitude can be seen to improve the efficacy of all the strategies tested, but is most noticeable for ACR and PL. Higher stimulation amplitudes also seem to be particularly beneficial for those systems with larger *k*_diag_. ACR can be seen to perform better than CR in all cases. Figure 11 shows the ACR method to be consistently more effective than PL at higher *k*_diag_, although the difference is marginal. This is indicative of the aforementioned effects of second harmonic terms in (32). Taken altogether, we predict the efficacy of ACR to be similar to PL stimulation for those systems where *a*_0_ ≃ 0.

**Figure 11.**
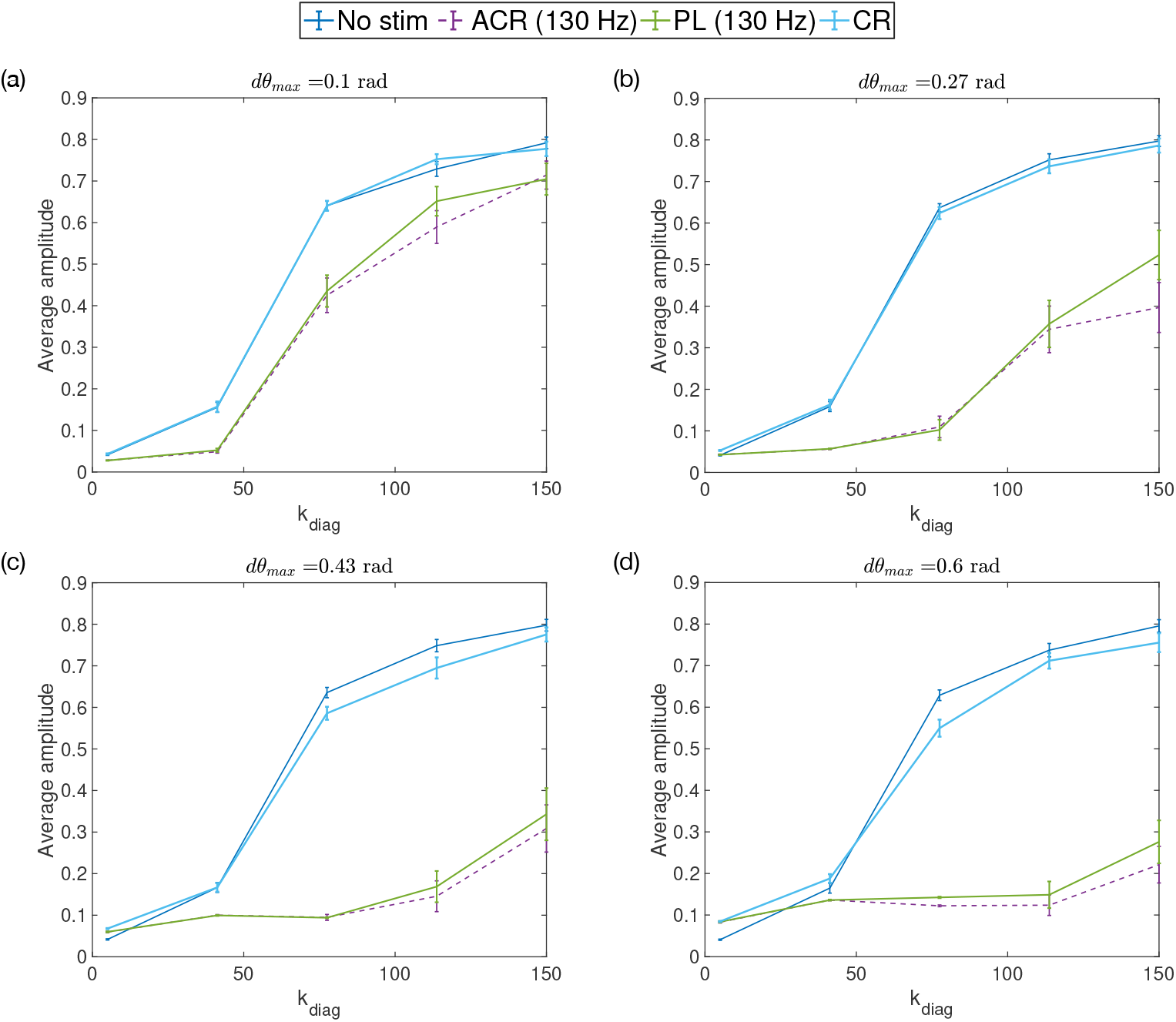
The average amplitude of a simulated Kuramoto system with a coupling constant *k*_diag_ for different stimulation strategies: no stimulation (no stim), Adaptive Coordinated Reset (ACR), phase-locked (PL) and Coordinated Reset (CR). During these simulations, the zeroth harmonic *a*_0_ of the nPRC *Z*(*θ*) was fixed to zero. Increasing *dθ*_max_ leading to larger stimulation amplitudes 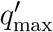 shows the effect of the second harmonic term in the amplitude response given by Equation (32).

## 6. Discussion

In this work we propose ACR as a method for DBS using multiple contacts. Unlike CR, the method is closed-loop and uses information about the system to determine when to apply stimulation. Using numerical simulation, we show that in many cases, substantial improvements to the efficacy can be achieved with the method. The mathematical description of ACR also predicts that the effectiveness of multi-contact stimulation is largely dependent on the form of the nPRC and in particular on the zeroth harmonic *a*_0_, which is related to whether it is type I or type II. We predict that for type II systems, where *|a*_0_*|* is small, stimulation on the basis of local quantities is unlikely to be beneficial. We also show that the dependency of the amplitude response on the local quantities of population *σ* becomes less at increasingly lower local amplitudes *ρ_σ_* but that the effects of stimulation are, in general, explicitly dependent on the state of the system and that providing stimulation without knowledge of this state is likely to be suboptimal. Following from this, it is worth discussing the feasibility of resolving this state in practice.

### 6.1. Practicalities of Using ICA

The ACR method assumes an underlying system of phase oscillators, which can be divided into small populations with the distribution of oscillators in each population satisfying the *ansatz* of Ott and Antonsen [26]. Equation (44) links the state to measurable quantities from the electrode and is of the form modelled by ICA [32]. The goal of ICA here is to resolve the *S* population quantities from *L* electrode measurements. Variations of the ICA problem exist depending on whether *S < L* (the overdetermined case), *S > L* (the underdetermined case) and *S* = *L* (the determined case). The determined case is perhaps the most common and more easily solved, since the mixing matrix *D* is invertible. However, we do not know the value of *S* a priori and therefore cannot know which ICA method is best suited. If we assume the case of *S* = *L*, then ICA will always resolve exactly *L* components. With this assumption, increasing the number of electrodes in a system has a definite purpose: it increases our potential to resolve the internal state. Assuming a larger number of populations also increases the validity of the small region approximation presented in Equation (38) and thus the accuracy of ACR. It may also be possible to obtain good approximations to the state by using *L < S* electrodes, since in some cases the weights *w_σ_* may be small for some populations and can hence be neglected. This together with the statistical nature of ICA, errors due to applying various signal processing techniques and noise within measurement would inevitably lead to some uncertainty when determining the population quantities. In addition to this, the amplitude response (32) is also dependent on the harmonics of *Z*(*θ*), which also need to be determined. Electrodes which record the population activity are also susceptible to recording the stimulation pulses themselves. This manifests in recordings as an artefact, which poses a challenge for closed-loop methods that rely on the real-time measurement of phases and amplitudes. Addressing the effects of stimulation artefacts is beyond the scope of this work, but we expect that significant suppression of the stimulation artefact would be required for ACR to be effective. This suppression may come as a byproduct of using ICA, which has been found by others [34, 35]. Alternatively, by recording through two contacts adjacent to a single stimulating contact, the properties of differential amplifiers can be used to suppress the stimulation artefact [36]. In this paper we have considered perturbations to neural populations using electrodes, but in principle, our theories should also be valid for other types of perturbation, such as optogenetic, where light pulses are used to perturb genetically modified neurons. This approach would eliminate the stimulation artefact and thus likely improve the real-world performance of ACR. In summary, the effectiveness of ACR in practice is likely to be dependent on both the ability of the model to capture the underlying dynamics of the system and our ability to resolve the state and parameters of the system. It is worth mentioning that the latter does not factor into the results presented in Section 5 since both the state and the parameters were taken directly from the simulation. Therefore the results we present for ACR (and PL) can be taken as an upper bound on performance.

### 6.2. Future Work

As a preliminary study, the simulations presented in this work provide a broad understanding of the potential efficacy and efficiency of ACR but there is scope for future work. Firstly, we could test the simulation on a greater variety of systems, changing both the distribution of natural frequencies, the noise parameter and the harmonics of *Z*(*θ*). Secondly, we could investigate how the constraints of the linear program (51) affect both the efficacy and efficiency of ACR. In addition to these, there is also considerable scope for understanding both the potential and effective real world performance of ACR, which could involve investigating the performance of ACR with the harmonics of *Z*(*θ*) estimated through machine learning, using the state variables obtained through ICA and finally testing ACR on patients.

## Appendix

## A. Model Fitting

## A.1. Feature Selection

In Section 3 we described how the similarity between two time series can be quantified using features extracted from the data. We define here a feature to be some transformation of the time series *F* (*t*) into a new function *y*(*ζ*), where *ζ* is in a new domain. For example, the PSD can be obtained by applying the Fourier transformation to the time series with *ζ* being the frequency in this case. We can then characterise a time series using a set of features. This set, though arbitrary, should be chosen so as to reproduce important properties of the data. For a set of *N_c_* features, we can construct a cost function *C*(**X**) for a vector of parameters **X** to be used in a local optimisation

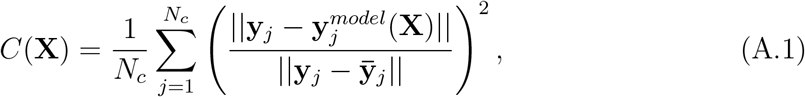

where **y** indicates a vector over the domain *ζ*. Here **y** and **y**^*model*^ are the features from the experimental and simulated data, respectively. The mean of an experimental feature is denoted by 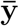. Each term in the summation is simply (1 − *R*^2^), where *R*^2^ is the standard coefficient of determination. Qualitatively, Equation (A.1) is simply the mean of (1 − *R*^2^) across all the features. Equation (A.1) quantifies the similarity between features obtained from simulated and experimental data. It can be seen that *C*(**X**) = 0 implies both sets of features are equal. When *C*(**X**) = 1, the fit of the model is no better than the mean 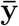. As previously mentioned in Section 3, the features reflecting the dynamics of the oscillations are chosen to be: the PSD, the PDF for the amplitude and the PSD of the envelope amplitude. We also use the averaged PRC as an additional feature to characterise the response of a particular patient.

**Figure A1.**
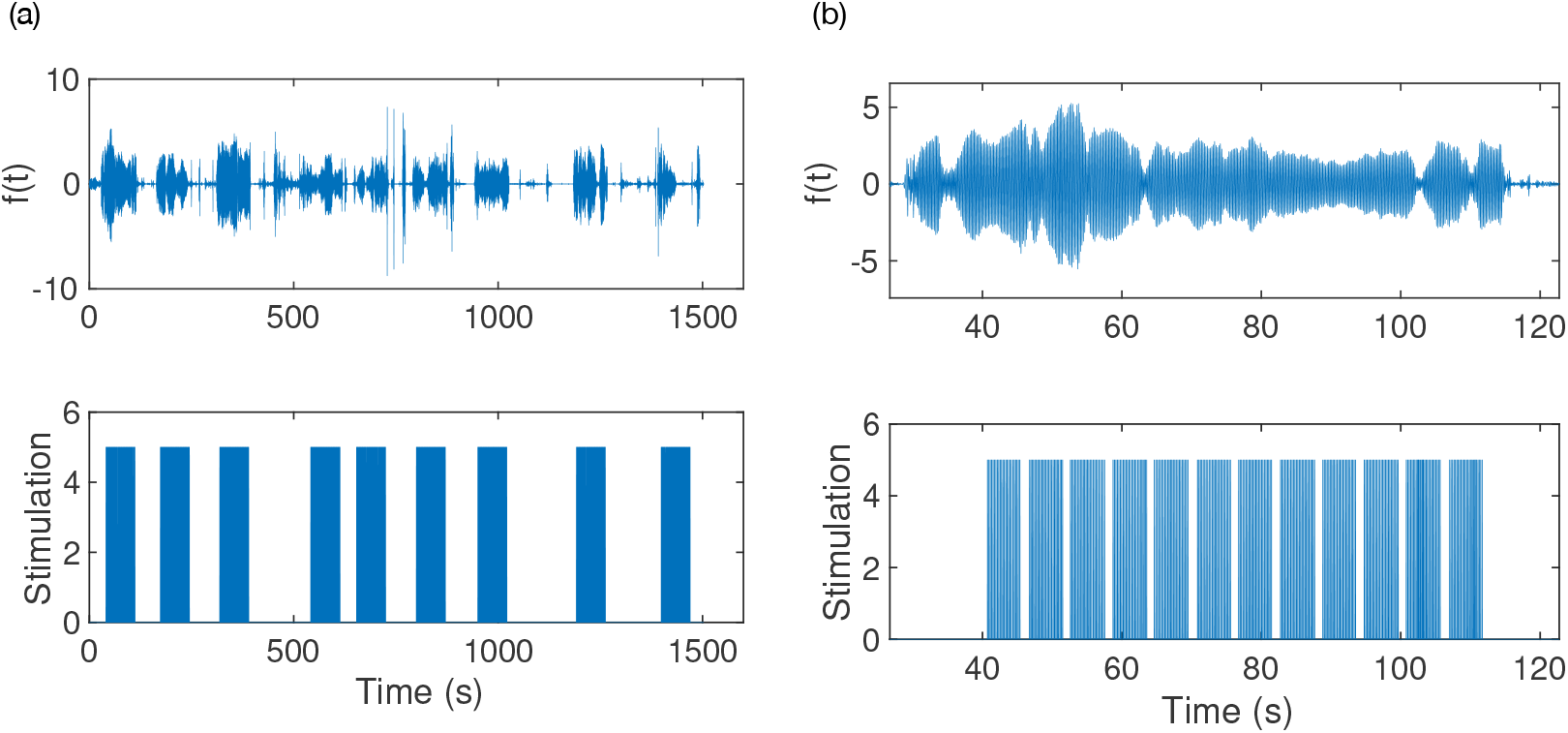
Experimental data for Patient 1 from the study of Cagnan et al [12]. Tremor oscillations are shown in the top panels. The bottom panels shows the stimulation triggers. (a) shows the entirety of the dataset consisting of stimulation provided over 9 trials. (b) shows a single trial which consists of 5 seconds of phase-locked stimulation over 12 phases.

## A.2. Experimental data

Cagnan et al [12] studied phase-locked DBS delivered according to the tremor in ET patients. Data was collected from 6 ET patients and 3 dystonic tremor patients. All patients gave their informed consent to take part in the study, which was approved by the local ethics committee in accordance with the Declaration of Helsinki. The data from this study can be obtained through an online repository [29].

Duchet et al [28] defined a criterion for assessing significance in the averaged ARCs and PRCs from the study of Cagnan et al. In their study, a patient is considered to have a significant response if both the ARC and PRC are found to be significant either according to an ANOVA test or cosine model F-test. Using this, they deemed 3 out of the 6 ET patients to have a significant response curve. We restrict our analysis to these 3 patients, who we shall refer to as patients 1, 5 and 6, as in the original study. The tremor data was filtered using a non-causal Butterworth filter of order 2 with cut-off frequencies at ±2 Hz around the tremor frequency. Stimulation was delivered over a set of trials (typically 9), with each trial consisting of 12 blocks of 5 second phase-locked stimulation at a randomly chosen phase from a set of 12. Each block of phase-locked stimulation was also separated by a 1 second interblock of no stimulation. The envelope amplitude and instantaneous phase were calculated using the Hilbert transform. As an example, the data for Patient 1 is shown in Figure A1. From this, the characteristics we identify as being desirable for our model to reproduce are: the frequency spectrum of the data, the bursts of oscillations and the sustained periods of low envelope amplitude. In addition to this we would also like the model to reproduce a given patient’s response to stimulation, as characterised by the averaged PRC.

**Table A1.** Parameters used when simulating the Kuramoto model for global optimisation.

| Parameter | Value | Description |
| --- | --- | --- |
| $\Delta t$ | 0.002 | Integration time step |
| $T$ | 5 | Stimulation time |
| $T_{ib}$ | 1 | Inter-block time |
| $T_{it}$ | 10 | Inter-trial time |
| $N_\sigma$ | 60 | Number of oscillators |
| $n_{\text{trials}}$ | 54 | Number of trials |

## A.3. Simulated data

For the *m*th local optimisation step, we simulate the Kuramoto model (18) using a parameter set **X**_*m*_ and obtain the feature set described in Section A.1. The stochasticity of the model naturally leads to variation in the features for a particular optimisation step. To stabilise this variation we average the features over *n*_trials_ = 54 trials. The simulation was configured to reproduce the methodology of Cagnan et al [12], namely that stimulation was delivered in blocks of trials, as described in Section A.2. A summary of the parameters used in the simulations is provided in Table A1.

Calculation of the cost function *C*(**X**) requires us to obtain the feature set for each instance of the simulation. The averaged PRC can be obtained according to the methods described in our previous paper [14] and elsewhere [12, 28]. The method is suitable for both experimental and simulated data, but can be computationally costly and generally unsuitable in an optimisation setting. To ensure computational feasibility, we use an approximation for the averaged PRC for part of the optimisation. A suitable approximation should be computationally cheap, stable and reasonably accurate. The requirement of stability precludes the use of the analytical expressions for the PRC (17), which are derived on the basis of an infinite system of oscillators satisfying the *ansatz* of Ott and Antonsen [26]. Situations affecting stability which may arise during optimisation include large values of the noise parameter 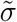, which may lead to a breakdown in the assumptions underlying (17). This motivates the need for alternative method, which we present here.

Assuming an infinite system of oscillators, the order parameter *r* can be written in integral form

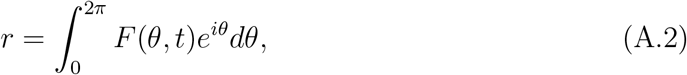

where *F* (*θ, t*) is the PDF for the oscillators. Differentiating with respect to time gives

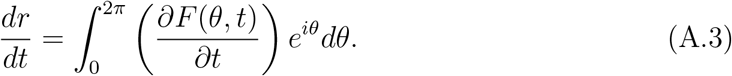

Using the stimulation part of (18), the continuity equation for *F* (*θ, t*) due only to stimulation can be written as

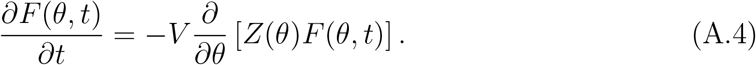

Inserting (A.4) into (A.3) gives

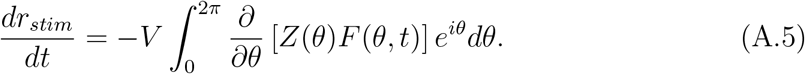

Using the polar form for 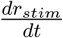 in (A.5) gives

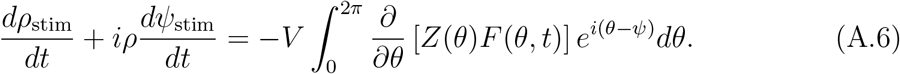

Expressions for the instantaneous ARC and PRC can be found by equating the real and complex parts of (A.6), respectively, leading to

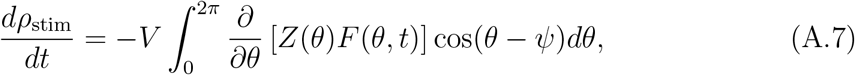

and

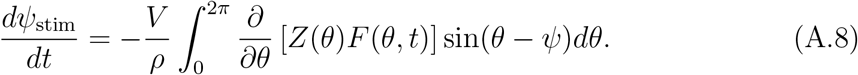

The averaged PRC can be expressed using a summation over the time points of stimulation {*t_m_*}

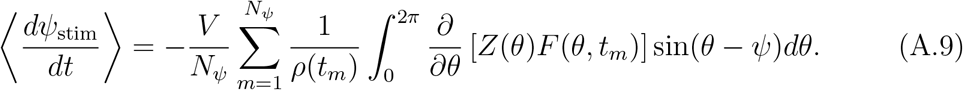

If we group these time points according to the phase *ψ*, with *N_ψ_* points for the phase *ψ*, then we can express the averaged response in terms of the PDFs conditioned on *ψ*

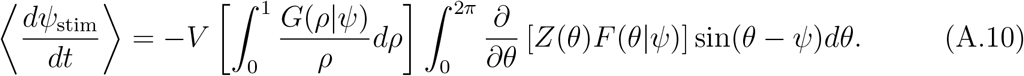

Equation (A.10) represents a computationally cheap way of estimating the averaged PRC since the PDF for the amplitude conditioned on the phase *G*(*ρ|ψ*) and the PDF for the oscillators *F* (*θ|ψ*) conditioned on the phase can be easily accumulated during simulation.

## A.4. Global optimisation

For each instance of the model output, we calculate the vector of features 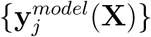 and their similarity with the experimental feature set {**y**_*j*_} measured by *C*(**X**). Since Equation (18) is a stochastic differential equation, the features 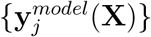 are averaged over a certain number of trials. The global minimum of the cost function *C*(**X**) corresponds to the set of optimisable parameters which best reproduces the experimental features. Starting at a configuration **X**_0_, we can reach a local minimum of *C*(**X**) using local optimisation. By repeating this process using many randomly generated starting configurations, a best fit can be obtained by taking the smallest local minimum. To obtain an initial configuration **X**_0_, we choose a value for each optimisable parameter by randomly sampling from a bounded uniform distribution. The bounds for each optimisable parameter are given in Table A2. Optimisation of the cost function *C*(**X**) was performed using custom written code in Matlab. The local optimisation was performed using the `fminsearch` function which uses the derivative-free Nelder-Mead simplex method of Lagarias et al [37].

Simultaneous optimisation of the Kuramoto parameters, together with those of the nPRC, is necessary to allow the model to fit to both the features reflecting the oscillation dynamics and the averaged PRC. In order to be computationally feasible, we performed the optimisation in stages. The principle here is to use a cheaper calculation to push the local optimisation towards a local minimum and then gradually refine the optimisation using a higher quality calculation. First, we performed the optimisation without stimulation, only fitting to those features representing the dynamics. We then used the parameters from our best fit in a second optimisation, using the features representing the dynamics and the theoretical approximation to the averaged PRC, for computational efficiency. Finally, the parameters from this best fit were used in a final optimisation, where the averaged PRC feature was instead calculated using the experimental methodology.

The best fits found through optimisation are shown in Figures 1 (for the dynamics) and 2 (for the response). Instances of output for the fitted models are shown in Figure 2 together with experimental data included for comparison. The parameters found through optimisation are provided in Table 2.

## B. Implications for Single Contact DBS

In this subsection we will review our results in the context of single contact DBS. Specifically, we want to understand the feasibility of a closed-loop DBS strategy which uses a feedback signal from a single contact. In the case of a single electrode contact, the voltage can be expressed as a summation over population activities using Equation (44)

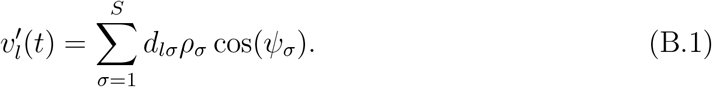

**Table A2.**
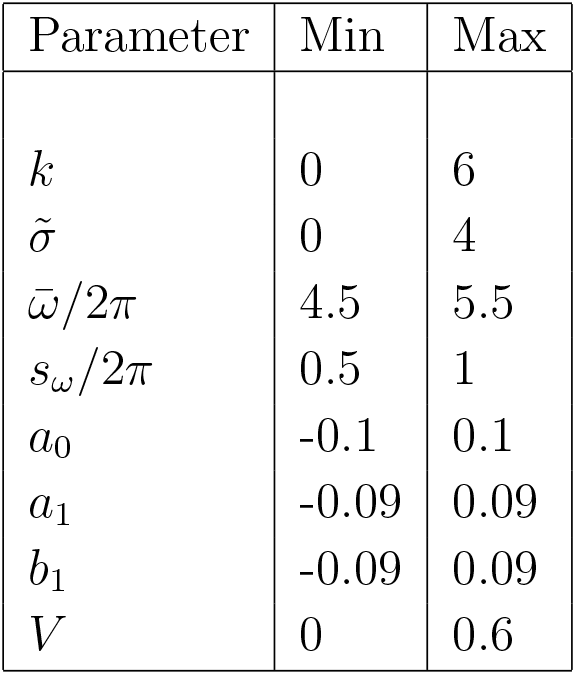
Bounds for the optimisable parameters used in generating random initial configurations. An initial configuration is generated by randomly sampling from a uniform distribution for each parameter within the bounds.

| Parameter | Min | Max |
| --- | --- | --- |
| $k$ | 0 | 6 |
| $\tilde{\sigma}$ | 0 | 4 |
| $\bar{\omega}/2\pi$ | 4.5 | 5.5 |
| $s_{\omega}/2\pi$ | 0.5 | 1 |
| $a_0$ | -0.1 | 0.1 |
| $a_1$ | -0.09 | 0.09 |
| $b_1$ | -0.09 | 0.09 |
| $V$ | 0 | 0.6 |

Comparing this with the expression for the global signal (24) and (25) (with *c* = 1 for simplicity) we can immediately see a correspondence between the matrix elements {*d_lσ_*} and the population weights {*w_σ_*}. The matrix elements encode the electrostatic properties of the medium and the electrode-population geometry. In theory, therefore, positioning the electrode has the effect of changing the matrix elements in the expansion given by Equation (B.1). An expression for the amplitude and phase of 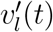 can be obtained using the analytic signal (1), namely

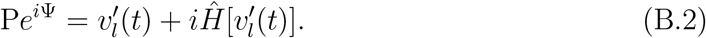

Then, inserting (B.1) into (B.2) and using the approximation (8) leads to

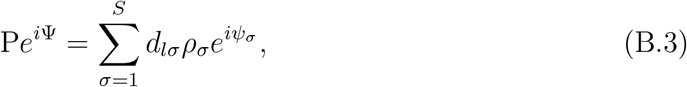

which has an identical form to (27). If the electrode is positioned such that the matrix elements coincide exactly with the population weights, although unlikely in general, then the amplitude P(*t*) would equal the synchrony *ρ*. In general, the electrode should be positioned so that P(*t*) is highly correlated to the symptom severity and hence *ρ*. Using (B.3), the derivation of the amplitude response due to stimulation can then proceed exactly as before, leading to an identical expression to (32) except with the population weights replaced with the matrix elements. Explicitly,

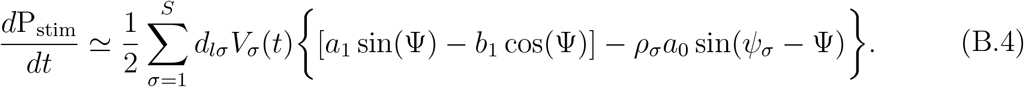

Since, by definition, P(*t*) should be correlated to symptom severity, it follows that Equation (B.4) can be used in a closed-loop DBS strategy. From this we also conclude that the effectiveness of single contact closed-loop DBS should also be dependent on *|a*_0_*|*.

In the cases where *|a*_0_*|* is non-negligible, knowledge of the population quantities *ρ_σ_* and *ψ_σ_* would be required for an effective closed-loop strategy. Therefore, by estimating *|a*_0_*|* for a particular system, we can go some way towards predicting the likely effectiveness of single contact closed-loop DBS.

